# Cross-Cancer Profiling of Cadherin-1 Reveals Context-Dependent Epithelial–Mesenchymal Transition Decoupling, Immune Heterogeneity, and Prognostic Variability in Epithelial Cancers

**DOI:** 10.64898/2026.05.22.727338

**Authors:** Md. Abdur Rahman, Sm Faysal Bellah, Md. Mustafizur Rahman

## Abstract

**Background:** CDH1 (E-cadherin) is a key epithelial adhesion molecule traditionally associated with tumor suppression and epithelial–mesenchymal transition (EMT). However, its roles across cancers remain incompletely understood, particularly within multilayer regulatory contexts involving genomic, epigenetic, transcriptional, and immune mechanisms.

**Methods:** CDH1 expression, survival associations, EMT-correlated gene profiles (VIM, SNAI1, ZEB1), immune infiltration patterns, immune checkpoint correlations (PDCD1, CD274, CTLA4), promoter methylation, and genomic alterations were assessed across five epithelial cancers, breast invasive carcinoma (BRCA), colon adenocarcinoma (COAD), lung adenocarcinoma (LUAD), ovarian cancer (OV), and stomach adenocarcinoma (STAD). Cross-platform validation was performed using TCGA/GDC datasets, GEPIA2, UALCAN, TIMER, KM Plotter, cBioPortal, and g:Profiler.

**Results:** CDH1 was overexpressed but showed variable prognostic significance; higher expression predicted better survival in COAD, LUAD and STAD, worse survival in BRCA and had no impact in OV. Classic inverse relationships between CDH1 and VIM or ZEB1 were evident only in STAD, and SNAI1 showed no consistent association. Immune infiltration patterns were tumor-specific, ranging from cytotoxic T-cell dominance in LUAD to macrophage-rich profiles in OV; immune checkpoint correlations were similarly context-dependent. Co-expressed genes were enriched for endomembrane transport rather than adhesion pathways. Promoter methylation patterns varied by cancer, whereas genomic alterations of CDH1 were rare.

**Conclusions:** CDH1 does not function as a universal epithelial or EMT marker across epithelial cancers. Instead, its associations with EMT, immune contexture, methylation, and prognosis are context-dependent, supporting a model of CDH1 as a heterogeneous regulator of epithelial plasticity. These findings challenge single-function interpretations and support cancer-specific CDH1 evaluation in translational research.

## 1 Introduction

Cadherin 1 (CDH1), which encodes E-cadherin, is a calcium-dependent adhesion molecule that maintains epithelial cell–cell junctions and tissue architecture<u>^1,2^</u>. E-cadherin stabilizes adherens junctions, preserves epithelial polarity, and restricts cellular motility through interactions with catenin complexes and the actin cytoskeleton<u>^3,4^</u>. Disruption of CDH1 impairs intercellular adhesion and contributes to tumor initiation and progression<u>^5^</u>. E-cadherin is also a central regulator of epithelial plasticity and EMT, a biological program associated with tumor invasion and metastatic dissemination<u>^6,7^</u>. Classical EMT is characterized by suppression of epithelial markers such as E-cadherin together with acquisition of mesenchymal traits that enhance migration and invasion<u>^8^</u>. Consequently, loss of E-cadherin has long been considered a hallmark of EMT and metastatic progression<u>^9,10^</u>. Multiple transcriptional and epigenetic mechanisms regulate CDH1 expression, including promoter methylation, EMT-associated transcriptional repression, and post-transcriptional modulation<u>^8,9^</u>.

However, emerging evidence indicates that EMT is not a strictly binary process and that many tumors exist in hybrid epithelial–mesenchymal states that retain partial epithelial features while acquiring invasive capacity<u>^11^</u>. In several malignancies, E-cadherin expression is preserved despite aggressive clinical behavior, metastatic dissemination, or collective cell migration<u>^12,13^</u>. These observations challenge the conventional interpretation of CDH1 as a uniformly suppressive factor in cancer progression and suggest that its biological role may vary substantially according to tissue context, tumor microenvironment, and regulatory state<u>^11,14^</u>.

Substantial evidence links CDH1 dysregulation to tumor progression across multiple epithelial malignancies<u>^14,15^</u>. Nevertheless, important conceptual gaps remain in understanding how CDH1 functions across different cancer types. Although CDH1 is widely regarded as a canonical epithelial marker, the extent to which its expression consistently maintains inverse relationships with mesenchymal programs across tumors remains unclear<u>^11,16^</u>. Similarly, whether CDH1-associated epithelial states are uniformly linked to immune infiltration patterns, immune checkpoint regulation, epigenetic modulation, and patient prognosis has not been systematically evaluated across epithelial cancers<u>^9,17^</u>. Most previous studies have examined CDH1 within single-cancer settings, limiting cross-cancer interpretation of its biological and clinical significance<u>^18^</u>.

To address these limitations, we performed a multilayer cross-cancer analysis of CDH1 across five epithelial malignancies, including BRCA, COAD, LUAD, OV, and stomach STAD. We evaluated whether classical associations between CDH1, EMT-related programs, immune contexture, epigenetic regulation, and prognostic outcomes remain conserved across distinct epithelial tumors. By integrating transcriptomic, prognostic, immune, methylation, and genomic analyses using multiple independent platforms, we aimed to characterize the context-dependent heterogeneity of CDH1 and to determine whether CDH1 functions as a universal epithelial marker or a multilayer regulator of epithelial plasticity in cancer.

## 2 Materials and Methods

### 2.1 Study design

The study employed a multi-platform in silico framework to investigate the biological and prognostic significance of CDH1 across five epithelial malignancies: BRCA, COAD, LUAD, OV, and STAD. These cancer types were selected based on their epithelial origin, established involvement of CDH1 dysregulation in tumor progression<u>^19-23^</u>, and the availability of harmonized transcriptomic and clinical data from publicly accessible cancer genomics repositories, primarily The Cancer Genome Atlas (TCGA)<u>^24^</u> and Genotype-Tissue Expression (GTEx) project<u>^25^</u>.

A stepwise analytical workflow was implemented to assess CDH1 expression patterns, sex-specific differences, prognostic significance, promoter methylation status, tumor immune associations, immune checkpoint correlations, mutation profiles, and functional heterogeneity across epithelial cancers. Analyses were performed using multiple publicly available bioinformatics platforms together with direct TCGA/GDC-derived cohort analyses, with each approach selected according to the specific biological objective. All data used in this study were obtained from publicly accessible databases. No individual-level identifiable data were accessed; therefore, ethical approval and informed consent were not required. The analytical workflow, datasets, and corresponding platforms used in this study are summarized in Table 1.

**Table 1.**
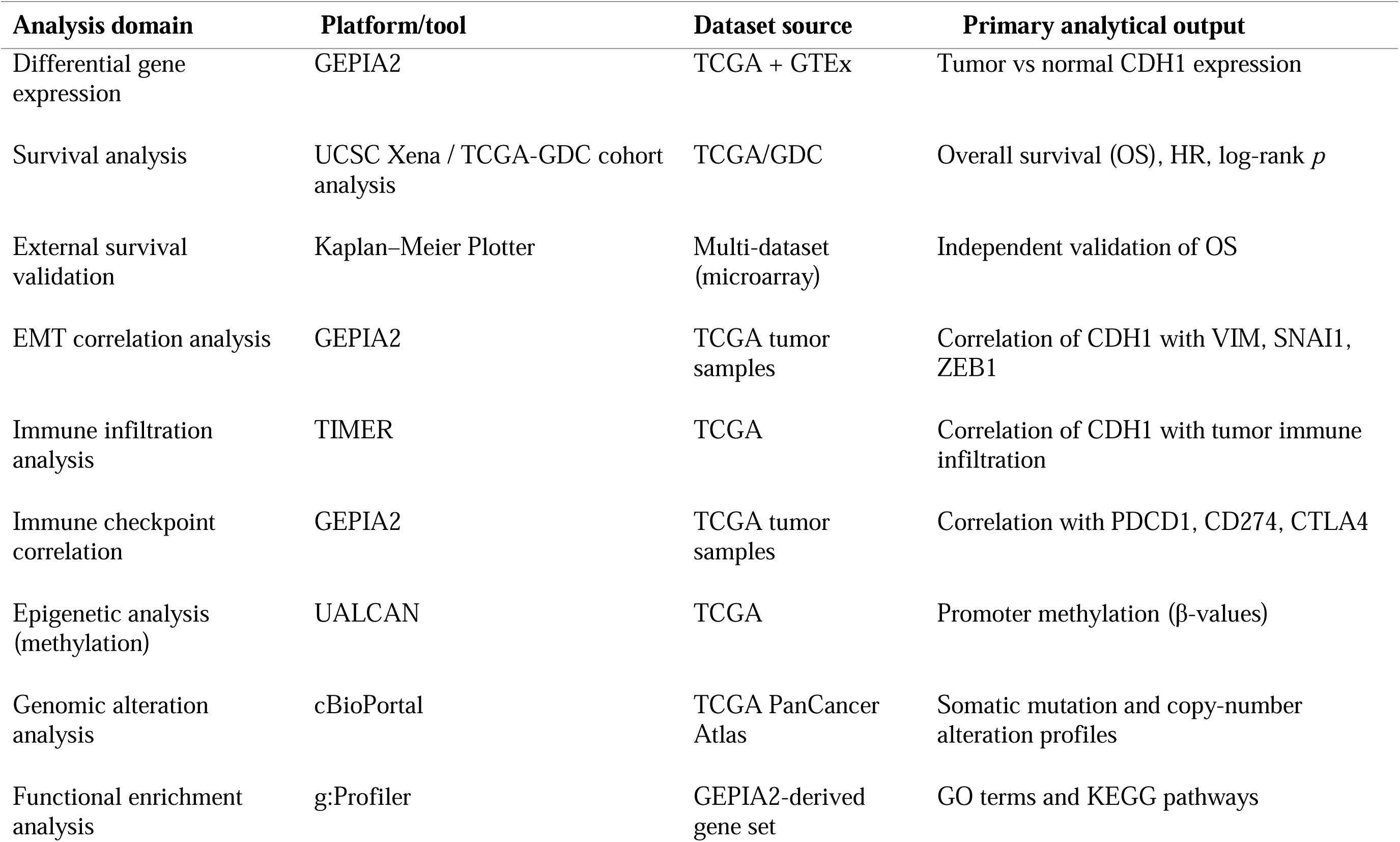
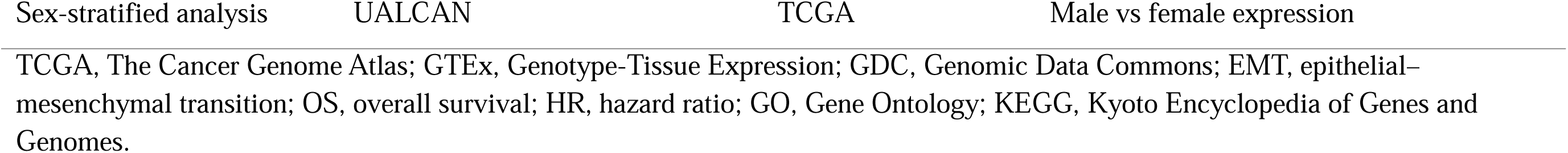
Analytical platforms, datasets, and outputs used in the study.

### 2.2 Differential gene expression analysis

CDH1 mRNA expression across tumor and normal tissues was evaluated using GEPIA2<u>^26^</u> and UALCAN<u>^27^</u>. GEPIA2 was used as the primary platform for differential expression analysis using integrated RNA sequencing data from TCGA and GTEx project<u>^24,25^</u>. Expression values were log2-transformed [log2(TPM + 1)]. The analysis was performed using the “Expression DIY–Box Plot” module with default parameters, including |log2 fold change (FC)| cutoff of 1, *p*-value cutoff of 0.01, and jitter size of 0.4. Matched normal tissues were derived from integrated TCGA normal and GTEx datasets within the GEPIA2 framework. Differential expression significance was calculated using the default statistical method implemented within GEPIA2, with disease state used as the grouping variable.

UALCAN was subsequently used to validate the expression patterns based on TCGA RNA-seq data. CDH1 transcription levels were compared between primary tumor and normal tissues, with statistical significance calculated using Student’s t-test implemented within the UALCAN platform.

### 2.3 Survival analysis

Overall survival (OS) analysis was primarily performed using TCGA/GDC-derived datasets obtained through the UCSC Xena platform<u>^28^</u>. STAR-TPM RNA sequencing expression matrices, phenotype files, and survival data were downloaded for BRCA, COAD, LUAD, OV, and STAD. CDH1 expression values were extracted using the Ensembl gene identifier ENSG00000039068. TCGA sample barcodes were converted to patient-level identifiers using the first 12 characters of each barcode, and expression data were matched with corresponding clinical survival information.

Overall survival time was measured in days using OS.time and OS status variables, where event status was coded as 1 for death and 0 for censored observations. Patients were stratified into high- and low-CDH1 expression groups according to the median expression value. Kaplan–Meier survival curves were generated, and differences between groups were evaluated using the log-rank test. Hazard ratios (HRs) and 95% confidence intervals (CIs) were estimated using univariate Cox proportional hazards regression analysis.

To validate TCGA-based findings, survival analysis was independently performed using the Kaplan–Meier Plotter platform, which integrates gene expression and survival data from multiple public repositories<u>^29^</u>. Patients were divided into high- and low-expression cohorts using the “auto-select best cutoff” option. Overall survival measured in months was selected as the survival endpoint, and probe selection was performed using the “JetSet best probe set” option with the Affymetrix probe ID 201131_s_at for CDH1 analysis<u>^30^</u>. Kaplan–Meier curves, hazard ratios, 95% confidence intervals, and log-rank *p*-values were extracted for comparison.

### 2.4 EMT correlation analysis

To evaluate whether canonical EMT-associated relationships with CDH1 were conserved across epithelial cancers, correlations between CDH1 expression and established EMT markers were assessed using GEPIA2<u>^26^</u>. The selected markers included VIM, representing a mesenchymal phenotype, and the EMT transcriptional regulators SNAI1 and ZEB1<u>^31^</u>. Analyses were performed across TCGA tumor samples using RNA-sequencing data integrated within GEPIA2. Gene expression values were log2-transformed [log2(TPM + 1)], and Pearson correlation coefficients were calculated to assess linear associations between CDH1 and EMT-related gene expression.

### 2.5 Immune infiltration analysis

The association between CDH1 expression and tumor immune infiltration was evaluated using TIMER3.0<u>^32^</u>. Correlation analysis was performed between CDH1 expression and the infiltration levels of major immune cell subsets, including CD4+ T cells, CD8+ T cells, macrophages, neutrophils, and dendritic cells. Spearman’s rank correlation coefficients were calculated using tumor purity-adjusted analyses implemented within the platform.

### 2.6 Immune checkpoint correlation analysis

To evaluate potential associations between CDH1 and immune checkpoint regulation, correlations were assessed between CDH1 expression and PDCD1 (PD-1), CD274 (PD-L1), and CTLA4 using GEPIA2<u>^26^</u>. Analyses were conducted across TCGA tumor samples using RNA-sequencing data integrated within the platform. Correlation analyses were performed using the GEPIA2 “Correlation Analysis” module. Gene expression values were log2-transformed [log2(TPM + 1)], and Pearson correlation coefficients were calculated.

### 2.7 Epigenetic and genomic alteration analysis

CDH1 promoter methylation was assessed using UALCAN, which provides TCGA DNA methylation data generated using the Illumina HumanMethylation450 platform<u>^27^</u>. Methylation levels were represented as β-values ranging from 0 (unmethylated) to 1 (fully methylated), derived from promoter-region CpG probes, including TSS200 and TSS1500. Tumor and normal tissues were compared using the unpaired two-sample Student’s t-test implemented within UALCAN. OV was excluded because the corresponding methylation data were unavailable.

Genomic alterations of CDH1 were analyzed using cBioPortal<u>^33,34^</u> across TCGA PanCancer Atlas cohorts<u>^35^</u>. Somatic mutations and copy-number alterations (CNAs) were evaluated using the Cancer Types Summary, OncoPrint, Mutations, and Plots modules. CNA data were based on GISTIC2.0 processing<u>^36^</u>. Alteration frequency, mutation type, mutation classification, and CNA categories were extracted. CDH1 mRNA expression was compared across CNA categories, including deep deletion, shallow deletion, diploid, gain, and amplification, to assess the association between copy-number variation and CDH1 mRNA expression.

### 2.8 Functional enrichment analysis

Genes positively correlated with CDH1 expression were identified using the GEPIA2 “Similar Genes Detection” module<u>^26^</u>. CDH1 was used as the query gene, and the top 100 positively correlated genes were selected from the combined BRCA, COAD, LUAD, OV, and STAD dataset.

Functional enrichment analysis was performed using g:Profiler<u>^37^</u>. Enrichment was assessed for Gene Ontology (GO) categories, including biological process (BP), molecular function (MF), and cellular component (CC), as well as Kyoto Encyclopedia of Genes and Genomes (KEGG) pathways. Statistical significance was determined using the g:SCS multiple testing correction method, with adjusted *p* < 0.05 considered significant.

### 2.9 Sex-stratified expression analysis

Sex-based differences in CDH1 expression were assessed using the UALCAN subgroup analysis module, which uses TCGA RNA-sequencing data with clinical annotations<u>^27,38^</u>. Analyses were performed for cancers with representation of both male and female patients, including COAD, LUAD, and STAD. BRCA and OV were excluded because of sex-specific cohort composition. CDH1 expression levels were compared between male and female groups within each eligible cancer type.

### 2.10 Statistical analysis

All statistical analyses were performed using the built-in functions of the respective analytical platforms. Statistical methods included Student’s t-test for group comparisons, platform-implemented differential expression analyses, Pearson or Spearman correlation analyses depending on the analytical platform, log-rank tests for survival comparisons, and univariate Cox proportional hazards regression for hazard ratio estimation. Where applicable, multiple testing correction was performed using the g:SCS method in g:Profiler. Statistical significance was defined as *p* < 0.05 unless otherwise specified. Key findings were cross-validated using independent analytical approaches, including GEPIA2, UALCAN, TCGA/GDC-derived cohort analyses, and Kaplan–Meier Plotter. Heatmap visualization and selected enrichment plots were generated using the SRplot online bioinformatics platform<u>^39^</u>.

## 3 Results

### 3.1 Differential expression of CDH1 across cancers

Differential expression of CDH1 was analyzed using GEPIA2 and independently validated using UALCAN across five epithelial malignancies. GEPIA2 analysis demonstrated significantly elevated CDH1 expression in tumor tissues compared with normal tissues in BRCA, COAD, LUAD, OV, and STAD (Fig. 1a), indicating broad but variable transcriptional upregulation across epithelial cancers.

**Figure 1.**
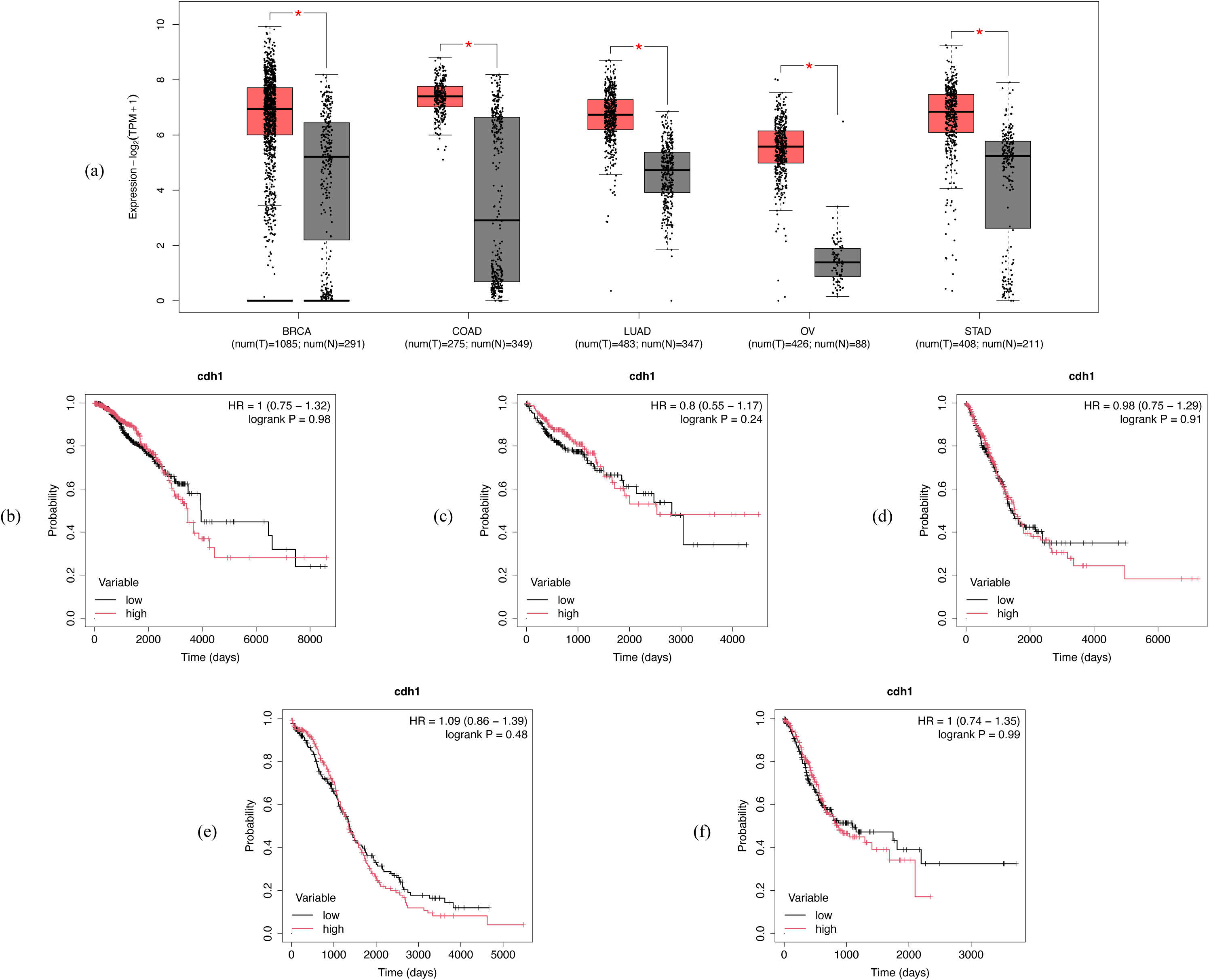
CDH1 expression and TCGA-based overall survival analysis across selected epithelial cancers. (a) Boxplot analysis comparing CDH1 mRNA expression between tumor and normal tissues in breast invasive carcinoma (BRCA), colon adenocarcinoma (COAD), lung adenocarcinoma (LUAD), ovarian serous cystadenocarcinoma (OV), and stomach adenocarcinoma (STAD). Expression values are presented as log2(TPM + 1). Red boxes represent tumor tissues and gray boxes represent normal tissues. Sample sizes for tumor (T) and normal (N) tissues are shown below each cancer type. Asterisks indicate statistically significant differences (*p* < 0.05). (b–f) Kaplan–Meier overall survival curves according to CDH1 expression in TCGA cohorts for (b) BRCA, (c) COAD, (d) LUAD, (e) OV, and (f) STAD. Patients were stratified into high- and low-expression groups based on CDH1 expression. Hazard ratios and log-rank p-values are shown within each survival plot. TPM, transcripts per million.

UALCAN validation largely supported these findings (Supplementary Fig. S1). CDH1 expression was significantly higher in BRCA, LUAD, and STAD tumor tissues compared with normal tissues (*p* < 0.001), consistent with GEPIA2 results. In contrast, COAD did not show a statistically significant difference between tumor and normal tissues (*p* = 0.092), with slight downregulation in tumors despite increased expression observed in GEPIA2, suggesting dataset-dependent variability. Due to the limited availability of normal ovarian tissue samples in TCGA, UALCAN-based tumor–normal comparison could not be performed for OV. However, stage-wise analysis demonstrated significant variation in CDH1 expression across ovarian cancer stages, particularly between stage II and stage III tumors (*p* = 0.023) (Supplementary Fig. S1d).

A cross-platform comparison of CDH1 expression patterns is summarized in Table 2. Overall, CDH1 expression tended to be elevated across most epithelial cancers, although the magnitude and statistical significance varied according to cancer type and analytical platform.

**Table 2.** Cross-platform validation of CDH1 expression.

| Cancer | GEPIA2<br>(Tumor vs Normal) | UALCAN<br>(Tumor vs Normal) | Interpretation |
| --- | --- | --- | --- |
| BRCA | Upregulated, * | Upregulated, $p < 0.001$ | Consistent |
| COAD | Upregulated, * | Downregulated (slight), $p = 0.092$ | Partial discordance |
| LUAD | Upregulated, * | Upregulated, $p < 0.001$ | Consistent |
| OV | Upregulated, * | Not available<sup>a</sup> | Limited validation |
| STAD | Upregulated, * | Upregulated, $p < 0.001$ | Consistent |

BRCA, breast invasive carcinoma; COAD, colon adenocarcinoma; GEPIA2, Gene Expression Profiling Interactive Analysis 2; LUAD, lung adenocarcinoma; OV, ovarian serous cystadenocarcinoma; STAD, stomach adenocarcinoma; TCGA, The Cancer Genome Atlas.
(\*) Asterisk indicates statistically significant differential expression in GEPIA2.
<sup>a</sup> UALCAN tumor-versus-normal comparison was not available for OV because normal ovarian tissue data were limited in TCGA.

### 3.2 Prognostic significance of CDH1

To evaluate the prognostic relevance of CDH1, overall survival analysis was primarily performed using TCGA/GDC-derived cohorts and independently validated using Kaplan–Meier Plotter.

TCGA-based survival analysis showed no significant association between CDH1 expression and overall survival in BRCA, COAD, LUAD, OV, or STAD (all *p* > 0.05). Median survival times were also comparable between CDH1-high and CDH1-low groups across all cohorts (Fig. 1(b-f)). Detailed TCGA-based overall survival statistics according to CDH1 expression are provided in Supplementary Table S1.

In contrast, Kaplan–Meier Plotter analysis revealed that elevated CDH1 expression was significantly associated with improved overall survival in COAD (HR = 0.73, *p* = 0.002), LUAD (HR = 0.74, *p* < 0.001), and STAD (HR = 0.74, *p* < 0.001), whereas no significant association was observed in OV (*p* = 0.12). A trend toward poorer survival was observed in BRCA (HR = 1.36, *p* = 0.003) (Supplementary Fig. S2).

Collectively, these findings indicate that the prognostic significance of CDH1 varies across epithelial malignancies and analytical platforms, supporting a context-dependent rather than universally conserved prognostic role.

### 3.3 EMT-associated correlations of CDH1 across epithelial cancers

To evaluate whether CDH1 expression maintained canonical EMT-associated relationships across epithelial cancers, correlation analyses were performed using GEPIA2 against the mesenchymal marker VIM and the EMT-associated transcription factors SNAI1 and ZEB1.

Significant inverse correlations between CDH1 and VIM were observed in BRCA (R = -0.20, *p* < 0.001) and STAD (R = -0.27, *p* < 0.001), indicating stronger associations with mesenchymal transition phenotypes in these cancers. A weaker inverse correlation was identified in COAD (R = -0.12, *p* = 0.045), whereas no significant association was observed in LUAD or OV. Correlation analyses involving SNAI1 demonstrated no significant associations across all analyzed cancers (all *p* > 0.05), suggesting limited involvement of SNAI1-mediated transcriptional repression in CDH1 regulation. In contrast, ZEB1 demonstrated substantial cancer-specific variation. A significant inverse correlation was identified exclusively in STAD (R = -0.25, *p* < 0.001), consistent with classical EMT regulation, whereas BRCA (R = 0.064, *p* = 0.034) and LUAD (R = 0.099, *p* = 0.030) showed weak positive correlations. COAD and OV demonstrated no significant association. A heatmap summarizing the correlations between CDH1 and EMT-associated genes across the analyzed cancers is presented in Fig. 2. The corresponding individual correlation plots and detailed Pearson correlation statistics for CDH1 and EMT-associated genes are provided in Supplementary Fig. S3 and Supplementary Table S2, respectively.

**Figure 2.**
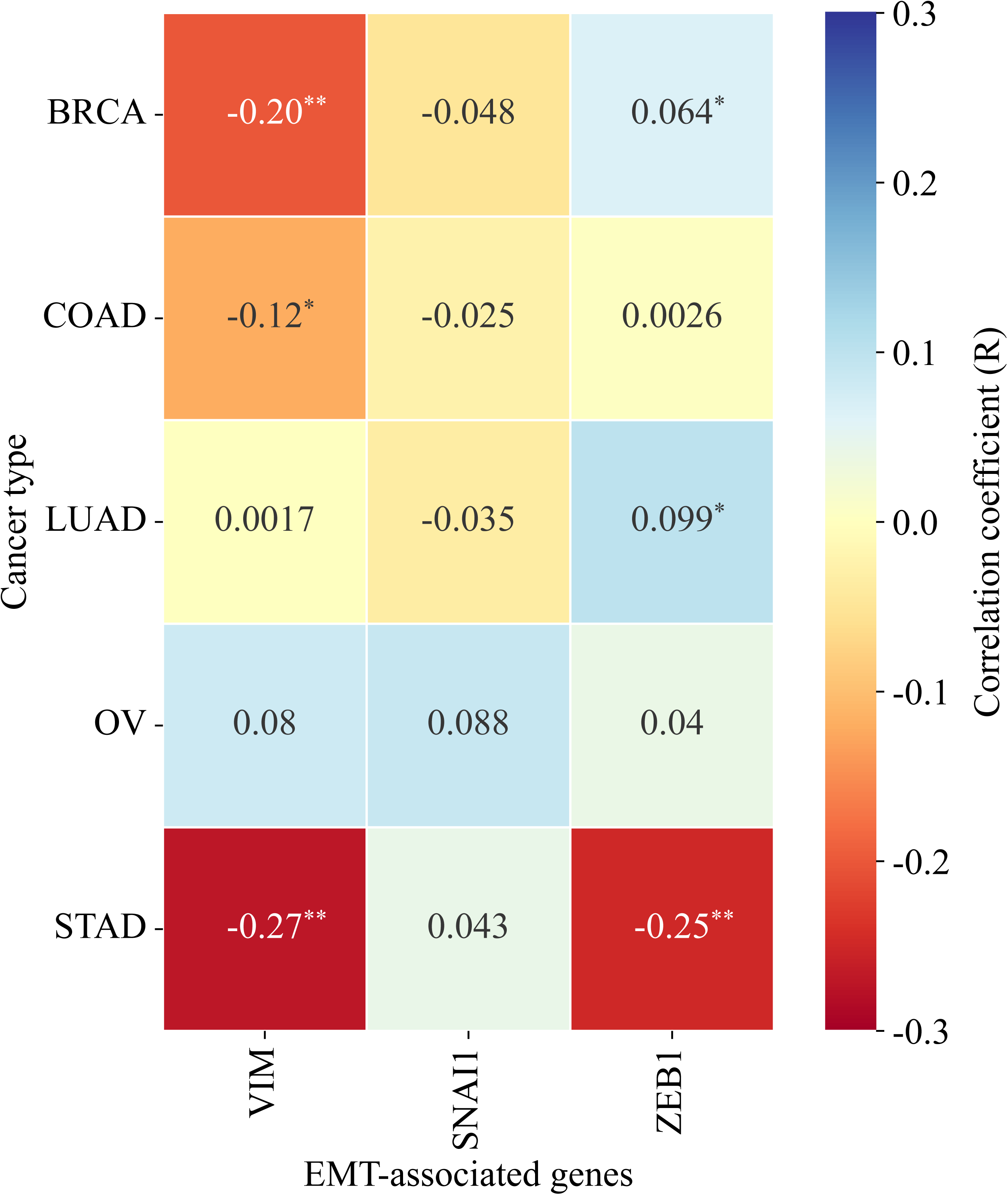
Correlation between CDH1 expression and EMT-associated genes across selected epithelial cancers. Heatmap showing Pearson correlation coefficients between CDH1 expression and EMT-associated genes VIM, SNAI1, and ZEB1 in breast invasive carcinoma (BRCA), colon adenocarcinoma (COAD), lung adenocarcinoma (LUAD), ovarian serous cystadenocarcinoma (OV), and stomach adenocarcinoma (STAD). Red indicates negative correlation and blue indicates positive correlation. Asterisks denote statistical significance: \**p* < 0.05; \*\**p* < 0.001. EMT, epithelial–mesenchymal transition.

Collectively, these findings indicate that CDH1-associated EMT relationships are not uniformly conserved across epithelial malignancies. Canonical inverse EMT-associated patterns were most evident in STAD, whereas BRCA demonstrated non-canonical EMT features, and COAD, LUAD, and OV exhibited weak or absent EMT-related associations despite elevated CDH1 expression.

### 3.4 Immune infiltration analysis

To explore potential immunological roles of CDH1, correlations between CDH1 expression and immune cell infiltration were evaluated using TIMER 3.0 across the five analyzed cancers.

Distinct cancer-specific immune infiltration patterns were observed (Fig. 3). In BRCA, CDH1 expression demonstrated positive associations with CD8+ T-cell (R = 0.104, *p* = 0.001) and macrophage infiltration (R = 0.128, *p* < 0.001), while inverse correlations were identified with CD4+ T cells (R = -0.240, *p* < 0.001) and dendritic cells (R = -0.136, *p* < 0.001), suggesting a complex immune regulatory profile. In COAD, CDH1 expression was positively associated with CD4+ T-cell (R = 0.145, *p* = 0.017) and macrophage infiltration (R = 0.152, *p* = 0.012), whereas no significant associations were observed for other immune cell subsets. LUAD demonstrated a more selective immune profile, with positive correlations identified for CD8+ T cells (R = 0.136, *p* = 0.003), macrophages (R = 0.099, *p* = 0.028), and dendritic cells (R = 0.150, *p* < 0.001). In OV, only macrophage infiltration demonstrated a significant positive association with CDH1 expression (R = 0.253, *p* < 0.001), whereas no significant correlations were observed for other immune cell populations. STAD demonstrated comparatively weaker immune associations, with only CD4+ T-cell infiltration showing a significant inverse correlation with CDH1 expression (R = -0.143, *p* = 0.006). The corresponding TIMER correlation plots and detailed immune infiltration statistics are provided in Supplementary Fig. S4 and Supplementary Table S3, respectively.

**Figure 3.**
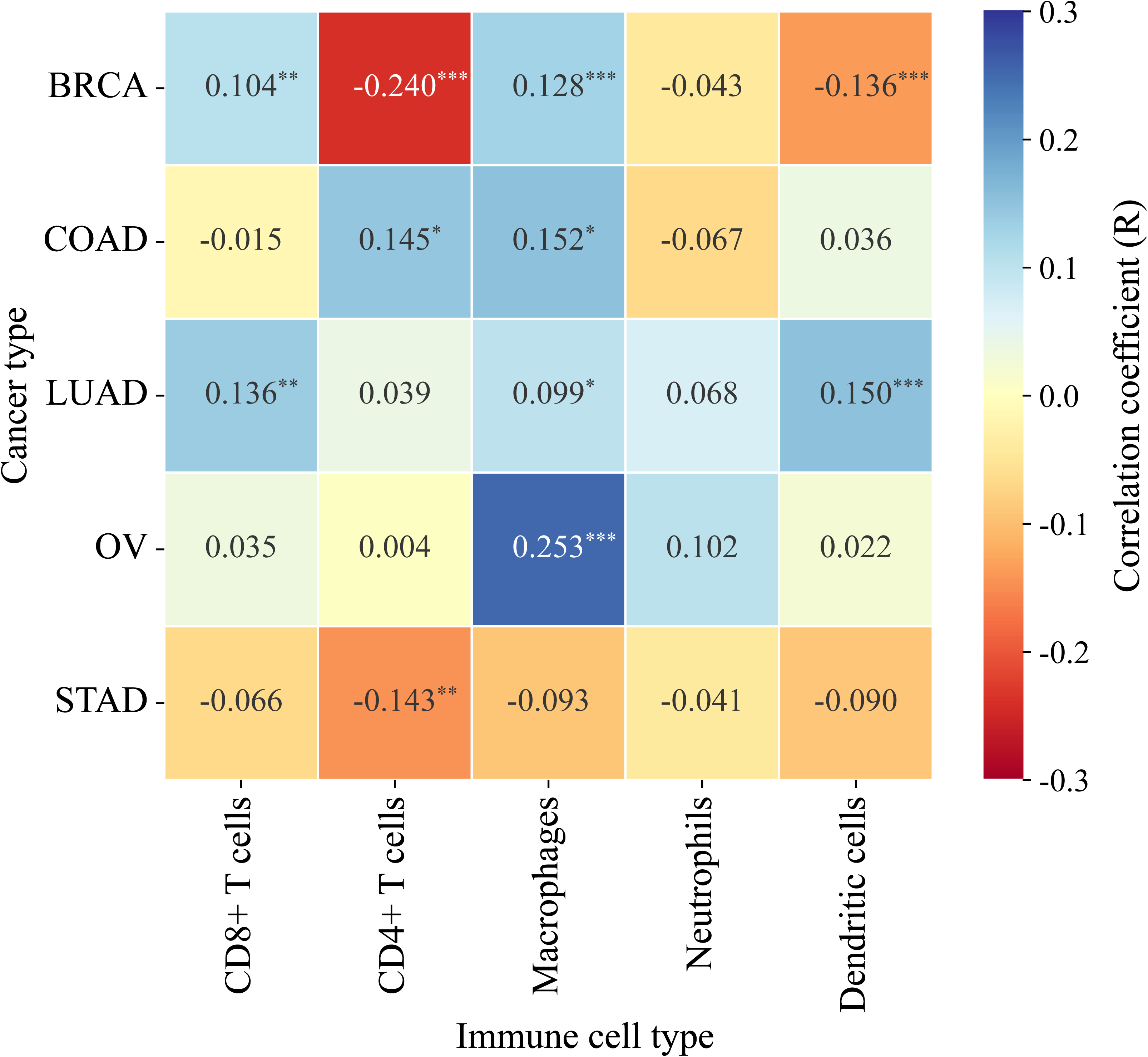
Correlation between CDH1 expression and immune-cell infiltration across selected epithelial cancers. Heatmap showing TIMER3.0-based purity-adjusted correlation coefficients between CDH1 expression and infiltration levels of CD8+ T cells, CD4+ T cells, macrophages, neutrophils, and dendritic cells in breast invasive carcinoma (BRCA), colon adenocarcinoma (COAD), lung adenocarcinoma (LUAD), ovarian serous cystadenocarcinoma (OV), and stomach adenocarcinoma (STAD). Red indicates negative correlation and blue indicates positive correlation. Asterisks denote statistical significance: \**p* < 0.05; \*\**p* < 0.01; \*\*\**p* < 0.001.

Overall, CDH1 demonstrated substantial immune heterogeneity across epithelial cancers. Macrophage infiltration showed the most consistent positive association, whereas CD4+ and CD8+ T-cell correlations varied according to tumor type. Neutrophils showed no significant associations across cancers. These findings suggest that CDH1-associated immune patterns are context-dependent and may reflect tumor-specific microenvironmental interactions rather than conserved immune regulatory mechanisms.

### 3.5 Association of CDH1 with immune checkpoint genes across cancers

Correlation analysis revealed distinct and cancer-specific patterns in the association between CDH1 expression and immune checkpoint genes.

In BRCA, CDH1 expression demonstrated weak but statistically significant negative correlations with PDCD1 (R = -0.16, *p* < 0.001), CD274 (R = -0.075, *p* = 0.014), and CTLA4 (R = -0.14, *p* < 0.001). These findings indicate a modest inverse association between CDH1 expression and immune checkpoint activity. In COAD, more consistent negative correlations were observed across all checkpoint markers, including PDCD1 (R = -0.21, *p* < 0.001), CD274 (R = -0.20, *p* < 0.001), and CTLA4 (R = -0.13, *p* = 0.033). This pattern suggests that higher CDH1 expression is associated with reduced immune checkpoint signaling in colorectal tumors.

In contrast, no significant associations between CDH1 expression and immune checkpoint genes were identified in LUAD or STAD. In OV, CDH1 expression demonstrated a weak but significant positive correlation with CD274 (R = 0.140, *p* = 0.004), whereas no significant associations were observed with PDCD1 or CTLA4 (Fig. 4). The corresponding individual GEPIA2 scatter plots are provided in Supplementary Fig. S5. Detailed Pearson correlation statistics are provided in Supplementary Table S4.

**Figure 4.**
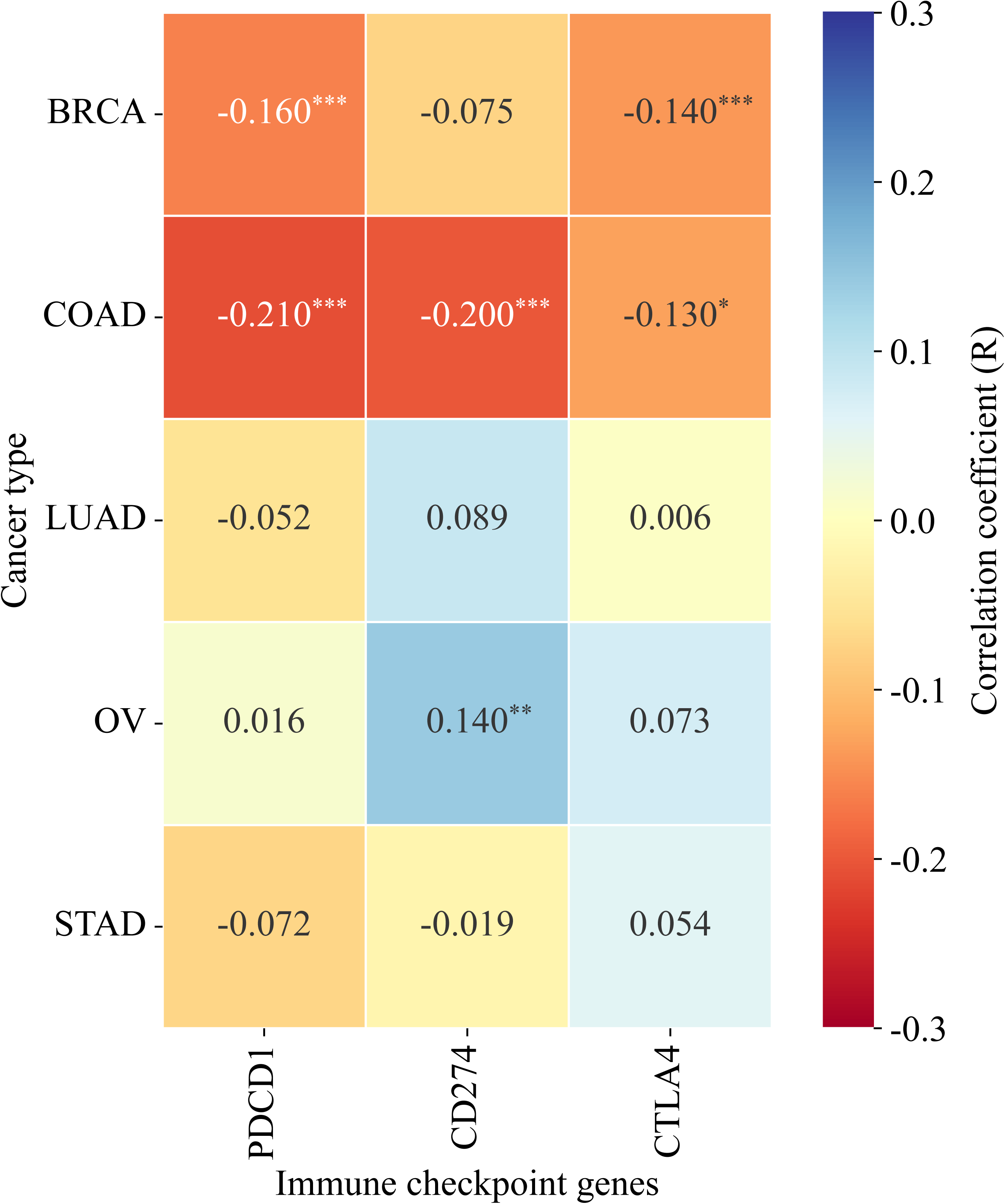
Correlation between CDH1 expression and immune checkpoint genes across selected epithelial cancers. Heatmap showing Pearson correlation coefficients between CDH1 expression and immune checkpoint genes PDCD1, CD274, and CTLA4 in breast invasive carcinoma (BRCA), colon adenocarcinoma (COAD), lung adenocarcinoma (LUAD), ovarian serous cystadenocarcinoma (OV), and stomach adenocarcinoma (STAD). PDCD1 encodes PD-1, CD274 encodes PD-L1, and CTLA4 encodes CTLA-4. Red indicates negative correlation and blue indicates positive correlation. Asterisks denote statistical significance: *p < 0.05; **p < 0.01; ***p < 0.001.

Overall, these findings demonstrate that the association between CDH1 and immune checkpoint regulation is highly heterogeneous across cancers. While CDH1 expression is linked to reduced checkpoint activity in COAD and, to a lesser extent, BRCA, it shows no consistent relationship in LUAD and STAD, and a distinct positive association with CD274 in OV. This variability highlights the context-dependent role of CDH1 in tumor immune modulation.

### 3.6 Genetic and epigenetic alterations of CDH1 across cancers

To investigate the regulatory basis of CDH1 dysregulation, promoter methylation and genomic alteration, analyses were performed across TCGA cohorts using UALCAN and cBioPortal, respectively.

Promoter methylation analysis revealed cancer-specific patterns (Supplementary Fig. S6). In BRCA, CDH1 promoter methylation levels remained low in both normal and tumor tissues, with only a modest difference between groups (*p* = 0.018). In COAD and STAD, tumor tissues demonstrated significantly reduced methylation compared with normal tissues (both *p* < 0.001), consistent with elevated CDH1 expression. In LUAD, a slight increase in promoter methylation was observed in tumor tissues (*p* = 0.028), although overall methylation levels remained low. Promoter methylation data were unavailable for OV.

Genomic alteration analysis revealed an overall low frequency of CDH1 alterations across cancers (Fig. 5), with relatively higher alteration rates observed in BRCA (13.65%) and STAD (10.68%), and lower frequencies in OV (3.77%), COAD (3.37%), and LUAD (1.41%). Mutations were the predominant alteration type and primarily consisted of truncating and splice-site variants, whereas copy-number alterations were comparatively infrequent.

**Figure 5.**
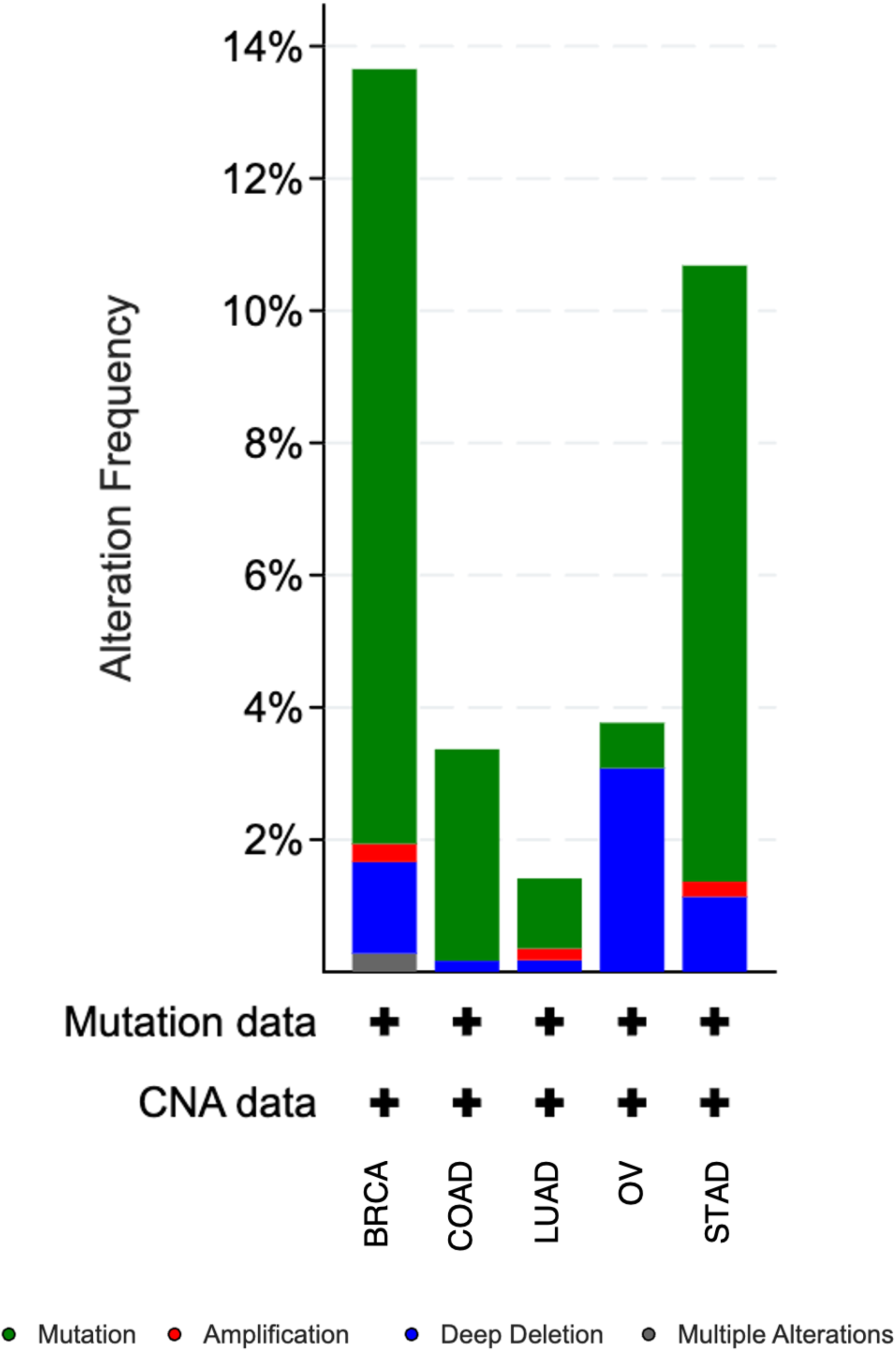
Genomic alteration landscape of CDH1 across selected epithelial cancers. cBioPortal analysis showing the frequency and type of CDH1 genomic alterations across selected TCGA PanCancer Atlas cohorts, including breast invasive carcinoma (BRCA), colon adenocarcinoma (COAD), lung adenocarcinoma (LUAD), ovarian serous cystadenocarcinoma (OV), and stomach adenocarcinoma (STAD). Alteration categories include mutation, amplification, deep deletion, and multiple alterations. Copy-number alteration and mutation data availability are indicated for each cancer cohort.

Copy-number analysis demonstrated a modest association between genomic dosage and CDH1 expression, with slightly higher expression observed in amplification and gain groups. However, substantial overlap across alteration categories suggests that copy-number changes are not the primary drivers of CDH1 expression variability. Detailed sample-level alteration patterns, mutation distribution, and copy-number-associated expression changes are shown in Supplementary Fig. S7, while cBioPortal-derived alteration frequencies are provided in Supplementary Table S5.

Collectively, these findings indicate that CDH1 dysregulation is not primarily driven by genomic alterations, but instead reflects a combination of epigenetic and non-genomic regulatory mechanisms that vary across cancer types.

### 3.7 Functional enrichment analysis

Functional enrichment analysis was performed to characterize biological processes associated with CDH1-correlated genes across the selected cancers. The input gene set was generated using the Similar Gene Detection module of GEPIA2 and analyzed using g:Profiler.

Enrichment analysis revealed significant clustering of CDH1-associated genes in processes related to intracellular transport, macromolecule localization, organelle organization, and endomembrane system organization. Notably, terms associated with vesicular trafficking, including ESCRT III complex disassembly, were also enriched (Supplementary Fig. S8). Cellular component analysis further supported this pattern, with significant enrichment observed in cytoplasmic ribonucleoprotein granules, Golgi apparatus subcompartments, cytosol, and endomembrane-associated structures. Notably, classical adhesion-related pathways, including adherens junction and cell–cell adhesion, were not among the dominant enriched terms.

Collectively, these findings suggest that CDH1-associated gene networks are preferentially linked to intracellular trafficking and membrane organization processes rather than canonical epithelial adhesion-related pathways.

### 3.8 Sex-stratified CDH1 expression analysis

Sex-stratified analyses were performed in COAD, LUAD, and STAD to evaluate whether sex influenced CDH1 expression patterns. BRCA and OV were excluded due to sex-specific cohort limitations.

In COAD, no significant differences in CDH1 expression were observed between male and female patients (*p* = 0.794), and tumor–normal comparisons within each sex were also not statistically significant. In contrast, LUAD demonstrated significantly elevated CDH1 expression in tumor tissues compared with normal controls in both male and female patients (*p* < 0.001), while no significant sex-based difference was observed (*p* = 0.777). Similarly, STAD demonstrated significantly increased CDH1 expression in tumor tissues in both sexes (*p* < 0.001), whereas no significant difference was identified between male and female patients (*p* = 0.122) (Supplementary Fig. S9 and Supplementary Table S6).

Overall, no significant sex-specific differences in CDH1 expression were observed across the analyzed cancers. These findings indicate that CDH1 expression patterns are primarily tumor-dependent rather than sex-dependent. Collectively, these findings demonstrate cancer-specific CDH1 profiles across expression, survival, EMT association, immune infiltration, immune checkpoint correlation, epigenetic regulation, and genomic alteration patterns (Table 3).

**Table 3.** Integrated summary of CDH1-associated findings across selected epithelial cancers.

| **Cancer** | **Expression** | **Survival** | **EMT association** | **Immune association** | **Checkpoint association** | **Epigenetic/genomic findings** | **Key message** |
| --- | --- | --- | --- | --- | --- | --- | --- |
| BRCA | Upregulated | No significant OS association | VIM-linked inverse association; no inverse SNAI1/ZEB1 pattern | Selective associations involving T cells, macrophages, and dendritic cells | Negative associations with PDCD1 and CTLA4 | Methylation assessed; highest alteration frequency | Upregulated CDH1 with non-classical EMT and selective immune remodeling |
| COAD | GEPIA2 upregulated; UALCAN non-significant | No significant OS association | Weak VIM-linked inverse association | CD4+ T-cell and macrophage associations | Negative associations with PDCD1, CD274, and CTLA4 | Methylation assessed; low-to-moderate alteration frequency | Platform-variable expression with limited EMT and immune-checkpoint associations |
| LUAD | Upregulated | No significant OS association | No VIM/SNAI1 association; weak positive ZEB1 association | CD8+ T-cell, macrophage, and dendritic-cell associations | No significant checkpoint association | Methylation assessed; lowest alteration frequency | Strong expression upregulation without clear EMT or prognostic coupling |
| OV | GEPIA2 upregulated; UALCAN tumor-normal unavailable | No significant OS association | No clear EMT association | Macrophage-associated immune pattern | CD274-specific positive association | Methylation unavailable; moderate alteration frequency | CDH1 upregulation with macrophage and CD274-linked immune features |
| STAD | Upregulated | No significant | Strong VIM- and ZEB1-linked | CD4+ T-cell negative | No significant checkpoint | Methylation assessed with | Most consistent classical EMT- |

| Cancer | Expression | Survival | EMT association | Immune association | Checkpoint association | Epigenetic/genomic findings | Key message |
| --- | --- | --- | --- | --- | --- | --- | --- |
|  |  | OS association | inverse association | association only | association | limited normal controls; high alteration frequency | linked CDH1 profile |

BRCA, breast invasive carcinoma; COAD, colon adenocarcinoma; EMT, epithelial–mesenchymal transition; GEPIA2, Gene Expression Profiling Interactive Analysis 2; LUAD, lung adenocarcinoma; OS, overall survival; OV, ovarian serous cystadenocarcinoma; STAD, stomach adenocarcinoma.

## 4 Discussion

In the present cross-cancer analysis, we found that CDH1 exhibits substantial context-dependent heterogeneity across epithelial malignancies rather than a uniformly conserved role. Although CDH1 encodes E-cadherin, a core adherens-junction protein that maintains cell–cell adhesion<u>^40^</u>, we observed that its behavior varied considerably among tumor types. Classical models position E-cadherin loss as a defining feature of EMT and metastatic progression<u>^7,41^</u>. However, our findings indicate that CDH1-associated EMT relationships, immune-associated correlations, and prognostic associations are not uniformly conserved across cancers. These observations support growing evidence that E-cadherin functions extend beyond a simple tumor suppressor model and instead reflect multilayer, context-dependent regulation associated with epithelial plasticity<u>^14,42^</u>.

In our study, we observed broad upregulation of CDH1 across multiple epithelial cancers, including BRCA, LUAD, STAD, and OV, with only partial variability in COAD. These results indicate that CDH1 expression is not uniformly suppressed during tumor progression and challenge the assumption that high CDH1 necessarily reflects a less aggressive phenotype<u>^43^</u>. Recent studies also suggest that CDH1 does not always decrease in cancer, as high or maintained expression has also been observed in aggressive and metastatic epithelial tumors<u>^44,45^</u>. Importantly, our survival analyses demonstrated marked prognostic heterogeneity across tumor types. Kaplan–Meier validation showed that high CDH1 correlated with favorable survival in COAD, LUAD, and STAD, but with poorer trends in BRCA and no significant effect in OV.

Similar context-dependent prognostic behavior was reported by Borcherding et al.<u>^46^</u>, who demonstrated that elevated E-cadherin expression may correlate with adverse clinical outcomes in specific breast cancer subtypes despite preservation of epithelial characteristics. Collectively, these findings suggest that CDH1 expression may not function as a universal prognostic biomarker and that its clinical significance is likely influenced by tumor-specific molecular characteristics and subtype heterogeneity.

In our EMT correlation analysis, we gained insight into the context-dependent behavior of CDH1 across epithelial cancers. Classical EMT models describe loss of E-cadherin as a defining event in epithelial–mesenchymal transition, typically accompanied by increased mesenchymal markers such as VIM and activation of EMT-associated transcription factors including SNAI1 and ZEB1<u>^31,47^</u>. However, our findings did not consistently follow this paradigm across tumor types. Significant inverse associations between CDH1 and VIM were primarily observed in BRCA and STAD, whereas weak or absent relationships were identified in LUAD, OV, and COAD. Similarly, SNAI1 did not demonstrate consistent inverse correlations with CDH1 despite its established role as a transcriptional repressor of E-cadherin<u>^47^</u>. In contrast, ZEB1 demonstrated a clearer inverse association predominantly in STAD, suggesting that ZEB1-mediated EMT regulation may be selectively preserved in specific tumor contexts<u>^48^</u>. These findings indicate that CDH1 expression alone may be insufficient to define EMT status across epithelial cancers. Emerging evidence suggests that EMT is better viewed as a dynamic spectrum rather than a binary process. In this framework, cells can occupy hybrid epithelial–mesenchymal states, retaining epithelial traits like E-cadherin while acquiring motility<u>^6^</u>. Such intermediate states may explain why elevated CDH1 expression did not uniformly correspond with less aggressive behavior or better survival in our cohort. Collectively, the observed heterogeneity in CDH1–EMT associations supports a model in which tumor-specific regulatory networks govern epithelial plasticity, extending beyond the traditional CDH1 suppression paradigm.

Our analysis of tumor immune infiltration and checkpoints further supports a context-dependent role for CDH1 beyond its classical epithelial functions. CDH1 showed heterogeneous associations with immune cells across cancers, indicating tumor-specific interactions with the microenvironment. For instance, macrophage infiltration was consistently positively correlated with CDH1, whereas correlations with T-cell populations varied by tumor type. In ovarian cancer (OV), CDH1 expression correlated more strongly with macrophage presence than with T cells, reflecting the known macrophage-rich, immunosuppressive microenvironment of OV<u>^49,50^</u>. In contrast, BRCA and LUAD demonstrated broader immune associations involving both macrophage and T-cell populations. Similar tumor-specific immune relationships involving CDH1 were previously reported by Fan et al.<u>^51^</u>, who observed negative associations between CDH1 expression and multiple immune-infiltrating cell populations in bladder cancer. Mechanistically, E-cadherin can engage the inhibitory receptor KLRG1 on T cells and natural killer (NK) cells, suggesting a direct link to immune regulation<u>^52^</u>. Consistent with these observations, we also identified heterogeneous correlations between CDH1 and immune checkpoint genes. Inverse correlations with PDCD1, CD274, and CTLA4 were observed in COAD and BRCA, whereas minimal associations were identified in LUAD and STAD. OV demonstrated a weak positive correlation between CDH1 and CD274, further supporting tumor-specific variation in checkpoint-associated expression patterns. Collectively, these findings suggest that the immunological relevance of CDH1 is highly cancer-type specific and may reflect distinct patterns of immune microenvironmental contexts across epithelial malignancies.

Our analysis also indicates that CDH1 dysregulation in epithelial cancers is not primarily driven by recurrent genomic alterations. We found that CDH1 mutation and copy-number alteration frequencies are relatively low across tumor types, implying that direct genetic inactivation contributes to CDH1 loss only in a subset of cancers. Although we did identify truncating and splice-site mutations, their limited prevalence is consistent with the view that non-genetic mechanisms predominate in CDH1 dysregulation across many epithelial malignancies<u>^53^</u>. Promoter methylation patterns also showed substantial variability between cancers. For example, CDH1 was relatively hypomethylated in COAD and STAD, consistent with its elevated expression, whereas LUAD displayed higher promoter methylation despite increased CDH1 expression. This mismatch suggests that DNA methylation alone does not fully dictate CDH1 transcription. Previous work has shown that CDH1 can be silenced by mutation or hypermethylation in cancers such as gastric cancer<u>^54,55^</u>. However, the modest methylation differences and low alteration frequencies identified in our study indicate that additional regulatory mechanisms, including transcription factor activity, post-transcriptional regulation, and tumor microenvironment-associated signaling, may substantially influence CDH1 expression across epithelial cancers.

Our enrichment analysis revealed that genes coexpressed with CDH1 are strongly associated with intracellular transport and vesicle-related processes, rather than classic cell–cell adhesion pathways. In other words, CDH1-linked networks in cancer appear tied to membrane organization and endomembrane systems. This is biologically meaningful because vesicle-mediated transport and membrane trafficking are increasingly recognized as important in tumor progression through receptor recycling, signaling, and cell–cell communication<u>^56,57^</u>. Therefore, the enrichment patterns of our study suggest that CDH1-associated pathways extend beyond maintaining epithelial integrity to include these broader regulatory processes. Previous studies have similarly proposed that E-cadherin-associated signaling may influence tumor biology through mechanisms beyond cell–cell adhesion, including modulation of intracellular signaling pathways<u>^43,46^</u>. The relative lack of enrichment in canonical adhesion pathways in our data supports the idea that CDH1’s transcriptional program in tumors encompasses complex intracellular and microenvironmental functions as well as its conventional structural role. In summary, these results imply that CDH1 might participate in endocytic or trafficking events (for instance, modulating receptor availability at the surface) that impact tumor behavior beyond simple junctional maintenance.

We evaluated CDH1 expression separately in male and female patients across TCGA and GTEx samples to determine if sex influences our results. Sex-stratified analyses showed largely comparable CDH1 expression patterns between male and female patients across the eligible cancers. In LUAD and STAD, tumor-associated CDH1 upregulation was observed in both sexes, whereas COAD showed no significant tumor-associated increase in either male or female patients. No significant male-versus-female differences were identified in any analyzed cancer. These findings suggest that the broader CDH1-associated patterns observed in our study are more likely driven by tumor-specific biology than by sex-related variation.

Collectively, these findings support a heterogeneous model of CDH1 biology across epithelial malignancies. Although CDH1 has traditionally been viewed as a suppressor of EMT and tumor progression, our analysis shows that elevated CDH1 expression can coexist with divergent survival outcomes, variable EMT-associated patterns, and distinct immune microenvironment interactions<u>^40^</u>. The retention or upregulation of CDH1 despite weak or inconsistent associations with classical EMT markers supports the emerging concept of epithelial plasticity and hybrid cellular states in tumor progression. Thus, the biological significance of CDH1 in cancer appears to extend beyond epithelial adhesion and may involve tumor-specific regulatory and microenvironmental functions.

The present study has several limitations. The analyses were primarily based on transcriptomic datasets, and mRNA expression may not fully reflect E-cadherin protein abundance, localization, or functional activity within tumors. The study also relied on retrospective public databases, which may contain variability related to cohort composition, clinical annotation, and tumor heterogeneity. Immune infiltration analyses were derived from computational estimation methods and therefore require experimental validation. In addition, detailed molecular subtype stratification and patient-level multi-omics integration analyses were not included, which may further refine the context-specific roles of CDH1 across cancers. Future experimental and clinically stratified studies will therefore be important to further evaluate the mechanistic and potential translational relevance of these findings.

## 5 Conclusion

This cross-cancer analysis demonstrates that CDH1 exhibits substantial context-dependent heterogeneity across epithelial malignancies. Although CDH1 is traditionally regarded as a canonical epithelial marker and regulator of epithelial–mesenchymal transition, its expression patterns, prognostic relevance, EMT associations, and immune-associated patterns varied markedly across tumor types. These findings indicate that CDH1 should not be interpreted uniformly as a surrogate marker of EMT or as a consistent prognostic indicator. Instead, its biological and clinical relevance appears to depend on tumor-specific molecular context, immune microenvironmental characteristics, and broader intracellular biological processes. The absence of conserved EMT-associated patterns across cancers highlights the limitations of applying classical EMT models universally. Overall, this study emphasizes the importance of integrative, multidimensional analyses when re-evaluating established cancer biomarkers. Future studies incorporating protein-level validation, molecular subtype stratification, and experimental models will be important to further clarify the mechanistic and clinical relevance of CDH1 across distinct cancer contexts.

## Acknowledgements

Not Applicable

## Author Contributions

**Md. Abdur Rahman**: Conceptualization; investigation; methodology; formal analysis; data curation; visualization; writing – original draft preparation; writing – review and editing. **Sm Faysal Bellah**: Methodology; validation; data interpretation; resources; writing – review and editing. **Md. Mustafizur Rahman**: Conceptualization; project administration, supervision, validation, data interpretation, writing – review and editing.

## Data Availability Statement

All datasets analyzed in this study are publicly available from TCGA/GDC and the respective online platforms referenced in the Methods section, including GEPIA2, UALCAN, TIMER, KM Plotter, cBioPortal, and g:Profiler.

## Funding Statement

The authors received no specific funding for this work.

## Conflict of Interest

The authors declare no conflicts of interest.

## Ethics Approval Statement

Ethical approval and informed consent were not required for this study because all analyses were performed using publicly available, de-identified datasets.

## Notes

### Competing Interest Statement

The authors have declared no competing interest.

## References

1. Takeichi M. The cadherins: cell-cell adhesion molecules controlling animal morphogenesis. Development. Apr 1988;102(4):639–55. doi:10.1242/dev.102.4.639

2. van Roy F, Berx G. The cell-cell adhesion molecule E-cadherin. Cell Mol Life Sci. Nov 2008;65(23):3756–88. doi:10.1007/s00018-008-8281-1

3. Ashkar F, Wu J. E-Cadherin and its signaling pathways: a novel target of dietary components in modulating cell migration and proliferation. Trends in Food Science & Technology. 2024;146:104398.

4. Le Bras GF, Taubenslag KJ, Andl CD. The regulation of cell-cell adhesion during epithelial-mesenchymal transition, motility and tumor progression. Cell Adh Migr. Jul-Aug 2012;6(4):365–73. doi:10.4161/cam.21326

5. Zou G, Huang Y, Zhang S, et al. E-cadherin loss drives diffuse-type gastric tumorigenesis via EZH2-mediated reprogramming. J Exp Med. Apr 1 2024;221(4):e20230561. doi:10.1084/jem.20230561

6. Nieto MA, Huang RY, Jackson RA, Thiery JP. Emt: 2016. Cell. Jun 30 2016;166(1):21-45. doi:10.1016/j.cell.2016.06.028

7. Thiery JP, Acloque H, Huang RY, Nieto MA. Epithelial-mesenchymal transitions in development and disease. Cell. Nov 25 2009;139(5):871–90. doi:10.1016/j.cell.2009.11.007

8. Nie F, Sun X, Sun J, Zhang J, Wang Y. Epithelial-mesenchymal transition in colorectal cancer metastasis and progression: molecular mechanisms and therapeutic strategies. Cell Death Discov. Jul 22 2025;11(1):336. doi:10.1038/s41420-025-02593-8

9. Xie Y, Wang X, Wang W, Pu N, Liu L. Epithelial-mesenchymal transition orchestrates tumor microenvironment: current perceptions and challenges. J Transl Med. Apr 2 2025;23(1):386. doi:10.1186/s12967-025-06422-5

10. Kalluri R, Weinberg RA. The basics of epithelial-mesenchymal transition. J Clin Invest. Jun 2009;119(6):1420–8. doi:10.1172/JCI39104

11. Jolly MK, Somarelli JA, Sheth M, et al. Hybrid epithelial/mesenchymal phenotypes promote metastasis and therapy resistance across carcinomas. Pharmacol Ther. Feb 2019;194:161–184. doi:10.1016/j.pharmthera.2018.09.007

12. Herndon ME, Ayers M, Gibson-Corley KN, et al. The highly metastatic 4T1 breast carcinoma model possesses features of a hybrid epithelial/mesenchymal phenotype. Dis Model Mech. Sep 1 2024;17(9):dmm050771. doi:10.1242/dmm.050771

13. Friedl P, Gilmour D. Collective cell migration in morphogenesis, regeneration and cancer. Nat Rev Mol Cell Biol. Jul 2009;10(7):445–57. doi:10.1038/nrm2720

14. Padmanaban V, Krol I, Suhail Y, et al. E-cadherin is required for metastasis in multiple models of breast cancer. Nature. Sep 2019;573(7774):439–444. doi:10.1038/s41586-019-1526-3

15. Berx G, Van Roy F. The E-cadherin/catenin complex: an important gatekeeper in breast cancer tumorigenesis and malignant progression. Breast Cancer Res. 2001;3(5):289–93. doi:10.1186/bcr309

16. Chae YK, Chang S, Ko T, et al. Epithelial-mesenchymal transition (EMT) signature is inversely associated with T-cell infiltration in non-small cell lung cancer (NSCLC). Sci Rep. Feb 13 2018;8(1):2918. doi:10.1038/s41598-018-21061-1

17. Thorsson V, Gibbs DL, Brown SD, et al. The Immune Landscape of Cancer. Immunity. Apr 17 2018;48(4):812–830 e14. doi:10.1016/j.immuni.2018.03.023

18. Samardali M, Samardaly J, Shanti I. A Comprehensive Literature Review of the CDH1 Mutation and Its Role in Gastric Cancer. Cureus. May 2025;17(5):e85072. doi:10.7759/cureus.85072

19. Pecina-Slaus N. Tumor suppressor gene E-cadherin and its role in normal and malignant cells. Cancer Cell Int. Oct 14 2003;3(1):17. doi:10.1186/1475-2867-3-17

20. Ryan CE, Fasaye GA, Gallanis AF, et al. Germline CDH1 Variants and Lifetime Cancer Risk. JAMA. Sep 3 2024;332(9):722–729. doi:10.1001/jama.2024.10852

21. Adib E, El Zarif T, Nassar AH, et al. CDH1 germline variants are enriched in patients with colorectal cancer, gastric cancer, and breast cancer. Br J Cancer. Mar 2022;126(5):797–803. doi:10.1038/s41416-021-01673-7

22. Juan W, Shan K, Na W, Rong-Miao Z, Yan L. The Associations of Genetic Variants in E-cadherin Gene With Clinical Outcome of Epithelial Ovarian Cancer. Int J Gynecol Cancer. Nov 2016;26(9):1601–1607. doi:10.1097/IGC.0000000000000829

23. Yu Q, Guo Q, Chen L, Liu S. Clinicopathological significance and potential drug targeting of CDH1 in lung cancer: a meta-analysis and literature review. Drug Des Devel Ther. 2015;9:2171–8. doi:10.2147/DDDT.S78537

24. 24. Cancer Genome Atlas Research N, Weinstein JN, Collisson EA, et al. The Cancer Genome Atlas Pan-Cancer analysis project. Nat Genet. Oct 2013;45(10):1113-20. doi:10.1038/ng.2764

25. Lonsdale J, Thomas J, Salvatore M, et al. The genotype-tissue expression (GTEx) project. Nature genetics. 2013;45(6):580–585.

26. Tang Z, Kang B, Li C, Chen T, Zhang Z. GEPIA2: an enhanced web server for large-scale expression profiling and interactive analysis. Nucleic Acids Res. Jul 2 2019;47(W1):W556–W560. doi:10.1093/nar/gkz430

27. Chandrashekar DS, Karthikeyan SK, Korla PK, et al. UALCAN: An update to the integrated cancer data analysis platform. Neoplasia. Mar 2022;25:18–27. doi:10.1016/j.neo.2022.01.001

28. Goldman MJ, Craft B, Hastie M, et al. Visualizing and interpreting cancer genomics data via the Xena platform. Nat Biotechnol. Jun 2020;38(6):675–678. doi:10.1038/s41587-020-0546-8

29. Gyorffy B, Lanczky A, Eklund AC, et al. An online survival analysis tool to rapidly assess the effect of 22,277 genes on breast cancer prognosis using microarray data of 1,809 patients. Breast Cancer Res Treat. Oct 2010;123(3):725–31. doi:10.1007/s10549-009-0674-9

30. Li Q, Birkbak NJ, Gyorffy B, Szallasi Z, Eklund AC. Jetset: selecting the optimal microarray probe set to represent a gene. BMC Bioinformatics. Dec 15 2011;12(1):474. doi:10.1186/1471-2105-12-474

31. Bakir B, Chiarella AM, Pitarresi JR, Rustgi AK. EMT, MET, Plasticity, and Tumor Metastasis. Trends Cell Biol. Oct 2020;30(10):764-776. doi:10.1016/j.tcb.2020.07.003

32. Cui H, Zhao G, Lu Y, et al. TIMER3: an enhanced resource for tumor immune analysis. Nucleic Acids Res. Jul 7 2025;53(W1):W534–W541. doi:10.1093/nar/gkaf388

33. Cerami E, Gao J, Dogrusoz U, et al. The cBio cancer genomics portal: an open platform for exploring multidimensional cancer genomics data. Cancer Discov. May 2012;2(5):401–4. doi:10.1158/2159-8290.CD-12-0095

34. Gao J, Aksoy BA, Dogrusoz U, et al. Integrative analysis of complex cancer genomics and clinical profiles using the cBioPortal. Sci Signal. Apr 2 2013;6(269):pl1. doi:10.1126/scisignal.2004088

35. Hoadley KA, Yau C, Hinoue T, et al. Cell-of-Origin Patterns Dominate the Molecular Classification of 10,000 Tumors from 33 Types of Cancer. Cell. Apr 5 2018;173(2):291–304 e6. doi:10.1016/j.cell.2018.03.022

36. Mermel CH, Schumacher SE, Hill B, Meyerson ML, Beroukhim R, Getz G. GISTIC2.0 facilitates sensitive and confident localization of the targets of focal somatic copy-number alteration in human cancers. Genome Biol. 2011;12(4):R41. doi:10.1186/gb-2011-12-4-r41

37. Kolberg L, Raudvere U, Kuzmin I, Adler P, Vilo J, Peterson H. g:Profiler-interoperable web service for functional enrichment analysis and gene identifier mapping (2023 update). Nucleic Acids Res. Jul 5 2023;51(W1):W207–W212. doi:10.1093/nar/gkad347

38. Chandrashekar DS, Bashel B, Balasubramanya SAH, et al. UALCAN: A Portal for Facilitating Tumor Subgroup Gene Expression and Survival Analyses. Neoplasia. Aug 2017;19(8):649–658. doi:10.1016/j.neo.2017.05.002

39. Tang D, Chen M, Huang X, et al. SRplot: A free online platform for data visualization and graphing. PloS one. 2023;18(11):e0294236.

40. Rubtsova SN, Zhitnyak IY, Gloushankova NA. Dual role of E-cadherin in cancer cells. Tissue Barriers. Oct 2 2022;10(4):2005420. doi:10.1080/21688370.2021.2005420

41. Pastushenko I, Blanpain C. EMT Transition States during Tumor Progression and Metastasis. Trends Cell Biol. Mar 2019;29(3):212–226. doi:10.1016/j.tcb.2018.12.001

42. Fang C, Kang Y. E-Cadherin: Context-Dependent Functions of a Quintessential Epithelial Marker in Metastasis. Cancer Res. Dec 1 2021;81(23):5800–5802. doi:10.1158/0008-5472.CAN-21-3302

43. Rodriguez FJ, Lewis-Tuffin LJ, Anastasiadis PZ. E-cadherin’s dark side: possible role in tumor progression. Biochim Biophys Acta. Aug 2012;1826(1):23–31. doi:10.1016/j.bbcan.2012.03.002

44. Elisha Y, Kalchenko V, Kuznetsov Y, Geiger B. Dual role of E-cadherin in the regulation of invasive collective migration of mammary carcinoma cells. Sci Rep. Mar 21 2018;8(1):4986. doi:10.1038/s41598-018-22940-3

45. Balamurugan K, Poria DK, Sehareen SW, et al. Stabilization of E-cadherin adhesions by COX-2/GSK3beta signaling is a targetable pathway in metastatic breast cancer. JCI Insight. Mar 22 2023;8(6):e156057. doi:10.1172/jci.insight.156057

46. Borcherding N, Cole K, Kluz P, et al. Re-Evaluating E-Cadherin and beta-Catenin: A Pan-Cancer Proteomic Approach with an Emphasis on Breast Cancer. Am J Pathol. Aug 2018;188(8):1910–1920. doi:10.1016/j.ajpath.2018.05.003

47. Dong B, Wu Y. Epigenetic Regulation and Post-Translational Modifications of SNAI1 in Cancer Metastasis. Int J Mol Sci. Oct 14 2021;22(20):11062. doi:10.3390/ijms222011062

48. Perez-Oquendo M, Gibbons DL. Regulation of ZEB1 Function and Molecular Associations in Tumor Progression and Metastasis. Cancers (Basel). Apr 7 2022;14(8):1864. doi:10.3390/cancers14081864

49. Lecker LSM, Berlato C, Maniati E, et al. TGFBI Production by Macrophages Contributes to an Immunosuppressive Microenvironment in Ovarian Cancer. Cancer Res. Nov 15 2021;81(22):5706–5719. doi:10.1158/0008-5472.CAN-21-0536

50. Colvin EK. Tumor-associated macrophages contribute to tumor progression in ovarian cancer. Front Oncol. 2014;4:137. doi:10.3389/fonc.2014.00137

51. Fan T, Xue L, Dong B, et al. CDH1 overexpression predicts bladder cancer from early stage and inversely correlates with immune infiltration. BMC Urol. Sep 21 2022;22(1):156. doi:10.1186/s12894-022-01103-7

52. Ito M, Maruyama T, Saito N, Koganei S, Yamamoto K, Matsumoto N. Killer cell lectin-like receptor G1 binds three members of the classical cadherin family to inhibit NK cell cytotoxicity. J Exp Med. Feb 20 2006;203(2):289–95. doi:10.1084/jem.20051986

53. Dongre A, Weinberg RA. New insights into the mechanisms of epithelial-mesenchymal transition and implications for cancer. Nat Rev Mol Cell Biol. Feb 2019;20(2):69–84. doi:10.1038/s41580-018-0080-4

54. Bayat M, Shirgir A, Kazemi Veisari A, Najjar Sadeghi R. Detection of CDH1 gene promoter hypermethylation in gastric cancer and chronic gastritis. Pract Lab Med. May 2024;40:e00406. doi:10.1016/j.plabm.2024.e00406

55. Graziano F, Arduini F, Ruzzo A, et al. Prognostic analysis of E-cadherin gene promoter hypermethylation in patients with surgically resected, node-positive, diffuse gastric cancer. Clin Cancer Res. Apr 15 2004;10(8):2784–9. doi:10.1158/1078-0432.ccr-03-0320

56. Mosesson Y, Mills GB, Yarden Y. Derailed endocytosis: an emerging feature of cancer. Nat Rev Cancer. Nov 2008;8(11):835–50. doi:10.1038/nrc2521

57. Caswell P, Norman J. Endocytic transport of integrins during cell migration and invasion. Trends Cell Biol. Jun 2008;18(6):257–63. doi:10.1016/j.tcb.2008.03.004

