## Supplementary Figure S1 for "Cross-Cancer Profiling of Cadherin-1 Reveals Context-Dependent Epithelial–Mesenchymal Transition Decoupling, Immune Heterogeneity, and Prognostic Variability in Epithelial Cancers"

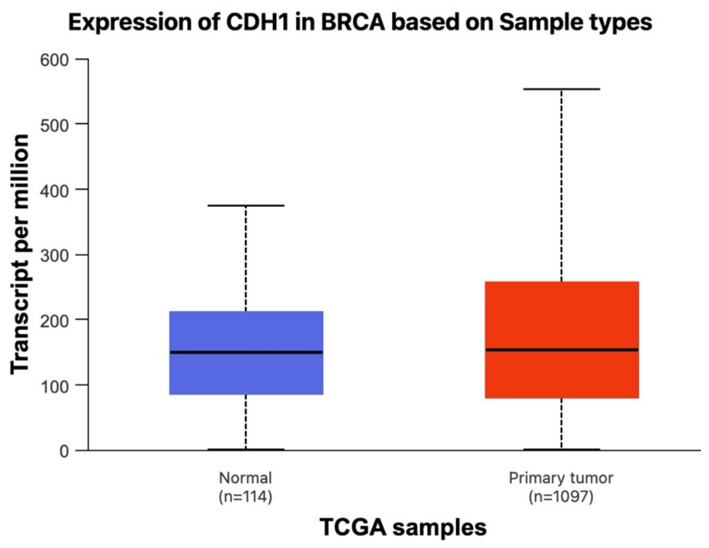


(a)

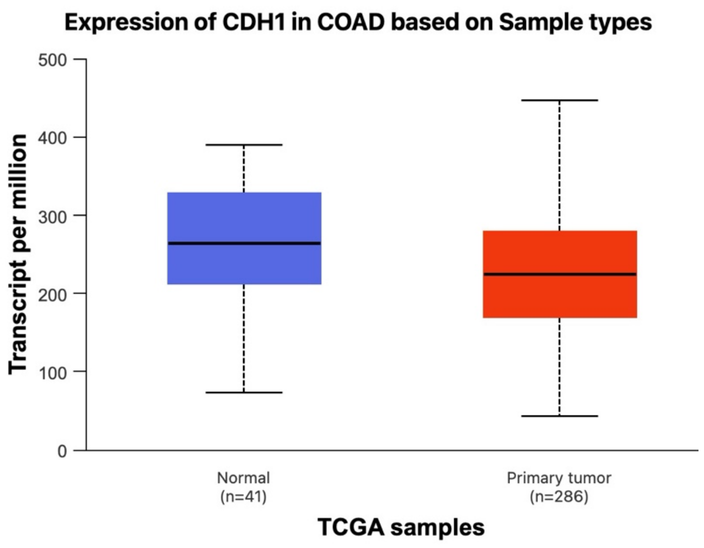


(b)

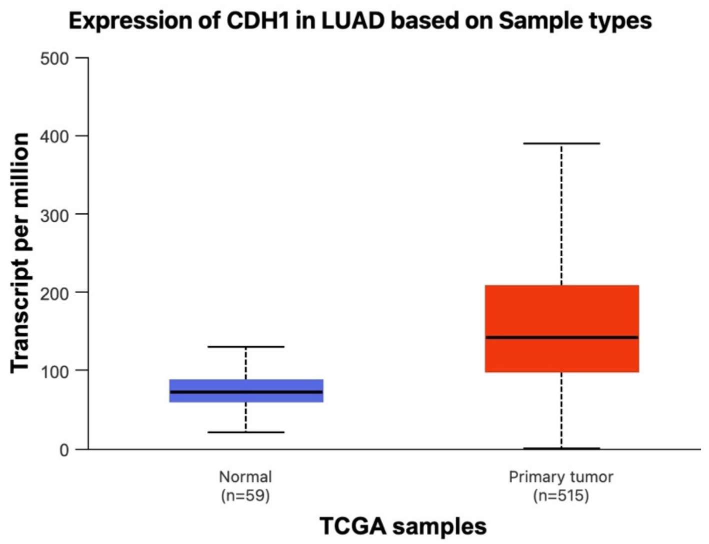


(c)

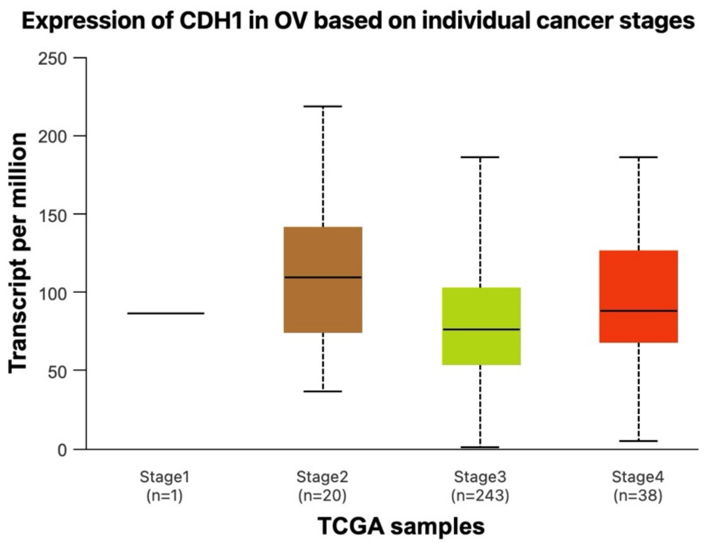


(d)

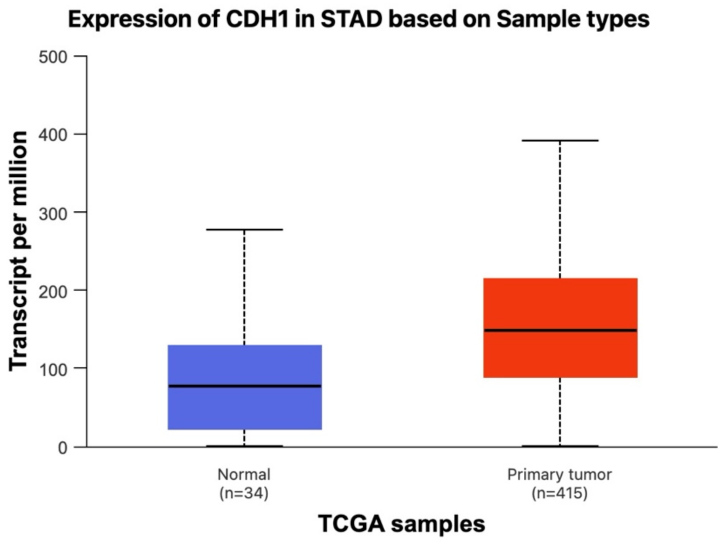


(e)

**Supplementary Figure S1. UALCAN validation of CDH1 expression in selected epithelial cancers.** CDH1 transcript expression was analyzed using UALCAN across **(a)** breast invasive carcinoma (BRCA), **(b)** colon adenocarcinoma (COAD), **(c)** lung adenocarcinoma (LUAD), **(d)** ovarian serous cystadenocarcinoma (OV), and **(e)** stomach adenocarcinoma (STAD). Normal-versus-primary tumor comparisons were available for BRCA, COAD, LUAD, and STAD. For OV, UALCAN provided stage-wise tumor expression analysis because normal ovarian tissue data were limited in TCGA.
