## Supplementary Figure S2 for "Cross-Cancer Profiling of Cadherin-1 Reveals Context-Dependent Epithelial–Mesenchymal Transition Decoupling, Immune Heterogeneity, and Prognostic Variability in Epithelial Cancers"

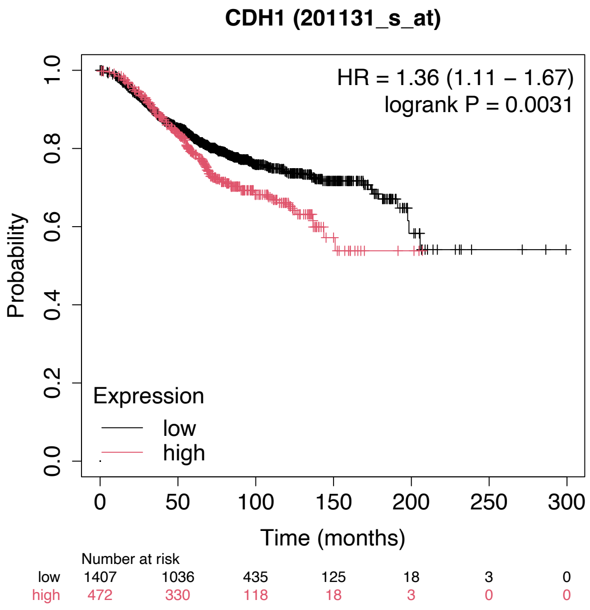


(a)

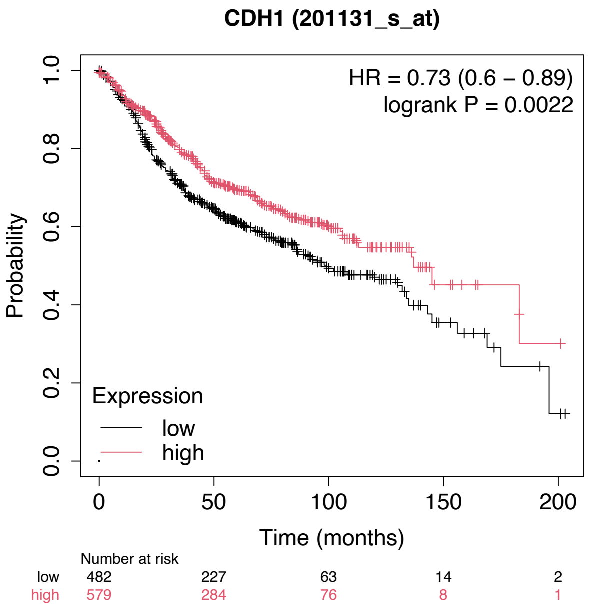


(b)

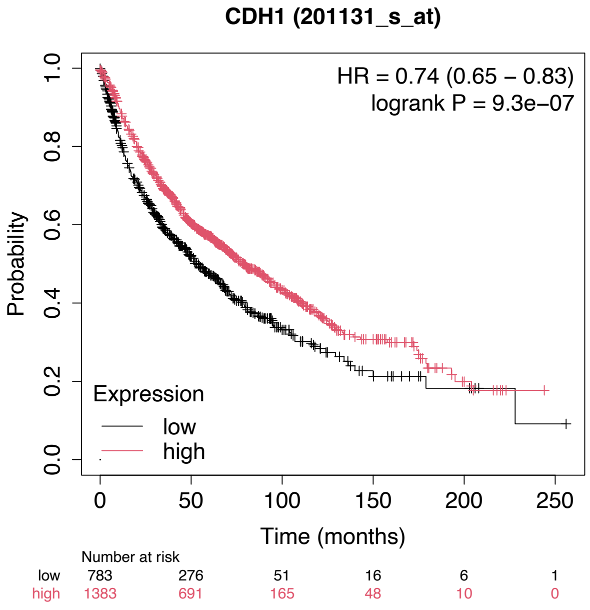


(c)

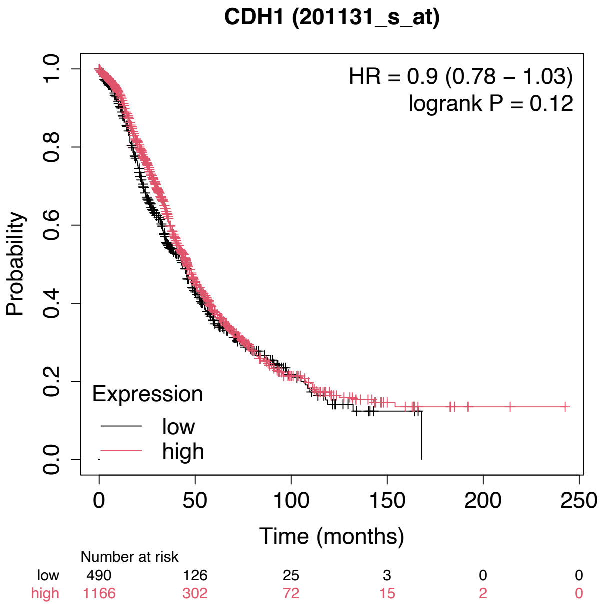


(d)

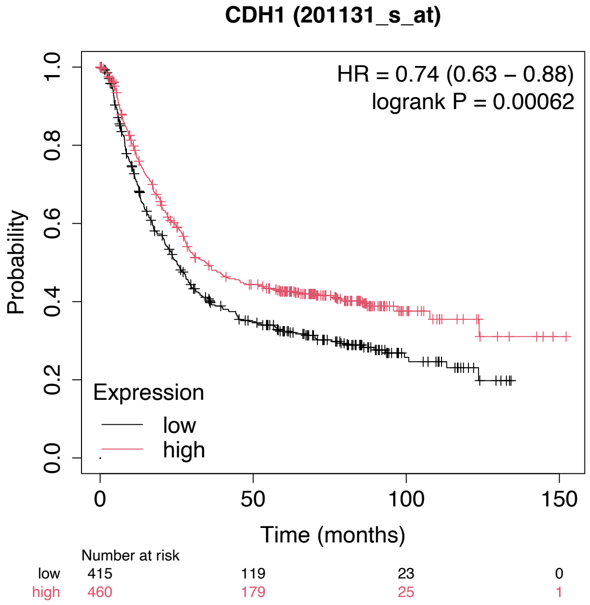


(e)

**Supplementary Figure S2. KM Plotter validation of the association between CDH1 expression and overall survival across selected epithelial cancers.** Kaplan–Meier survival curves generated using KM Plotter showing the association between CDH1 expression and overall survival in **(a)** breast invasive carcinoma (BRCA), **(b)** colon adenocarcinoma (COAD), **(c)** lung adenocarcinoma (LUAD), **(d)** ovarian serous cystadenocarcinoma (OV), and **(e)** stomach adenocarcinoma (STAD). Patients were stratified into high- and low-CDH1 expression groups according to the KM Plotter-defined cutoff. Hazard ratios, 95% confidence intervals, log-rank p-values, and numbers at risk are shown within the individual plots.
