## Supplementary Figure S3 for "Cross-Cancer Profiling of Cadherin-1 Reveals Context-Dependent Epithelial–Mesenchymal Transition Decoupling, Immune Heterogeneity, and Prognostic Variability in Epithelial Cancers"

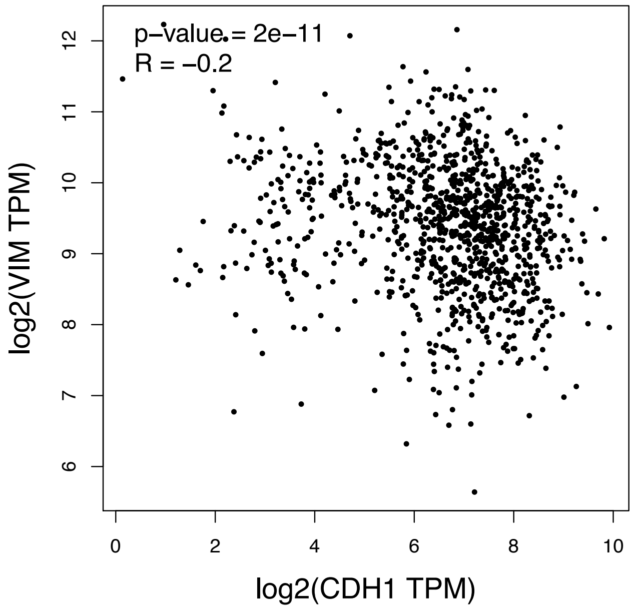


(a)

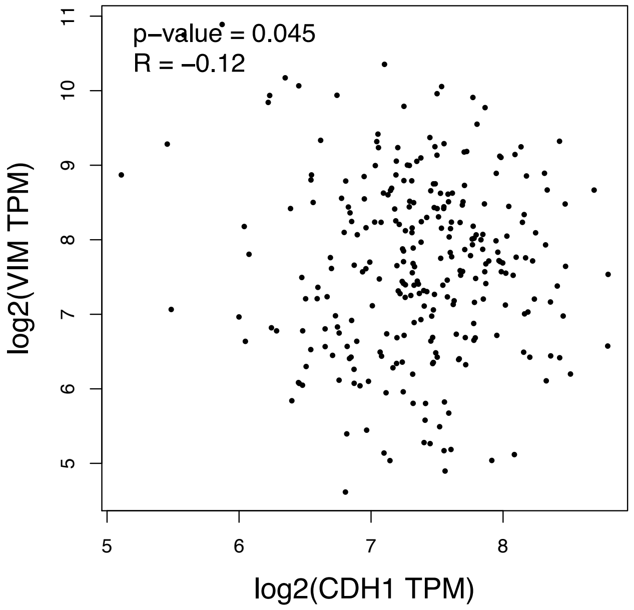


(b)

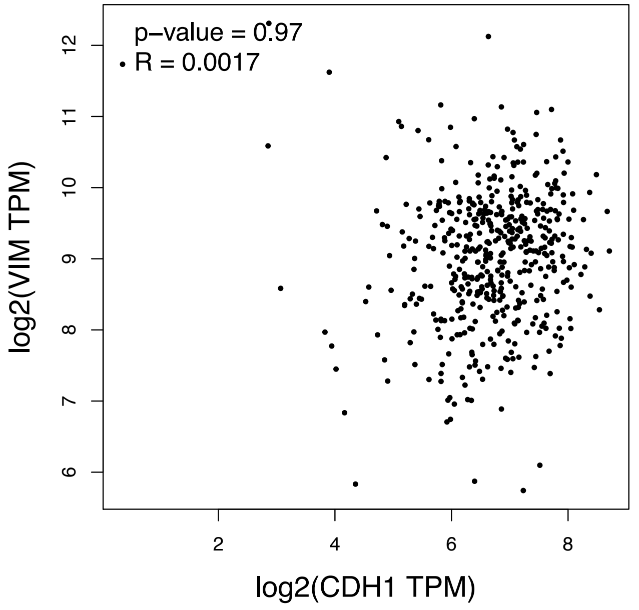


(c)

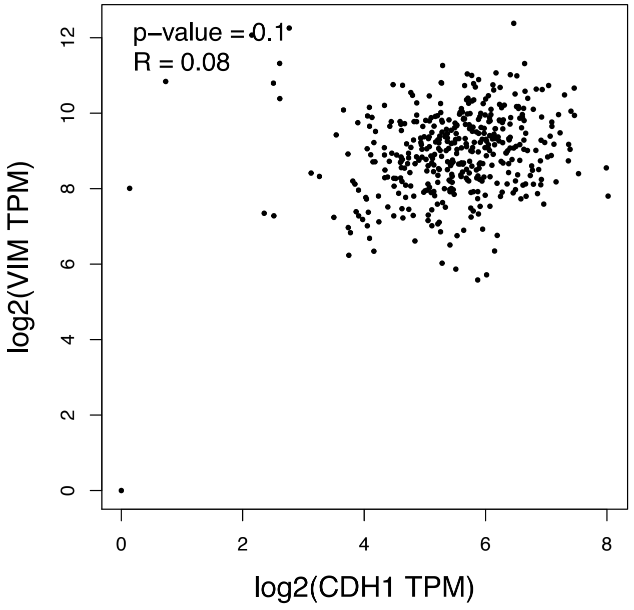


(d)

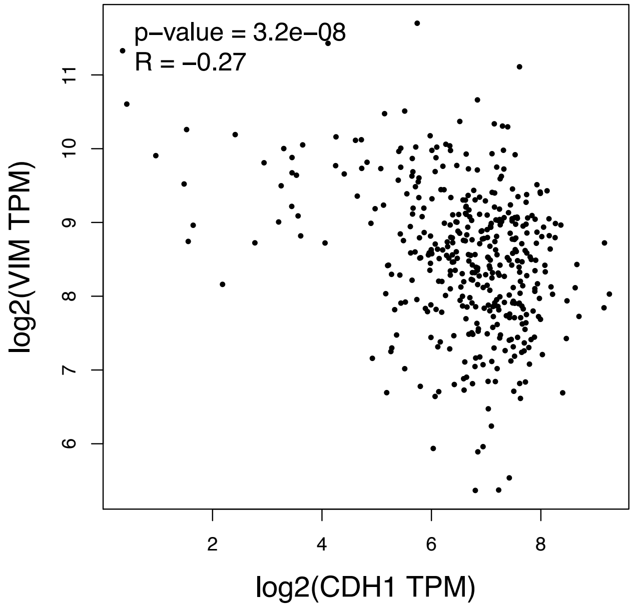


(e)

**Supplementary Figure S3A. Correlation between CDH1 and VIM expression across selected epithelial cancers.**
Pearson correlation analyses between CDH1 and VIM expression were performed using GEPIA2 across **(a)** breast invasive carcinoma (BRCA), **(b)** colon adenocarcinoma (COAD), **(c)** lung adenocarcinoma (LUAD), **(d)** ovarian serous cystadenocarcinoma (OV), and **(e)** stomach adenocarcinoma (STAD). Correlation coefficients and p-values are shown within each plot.

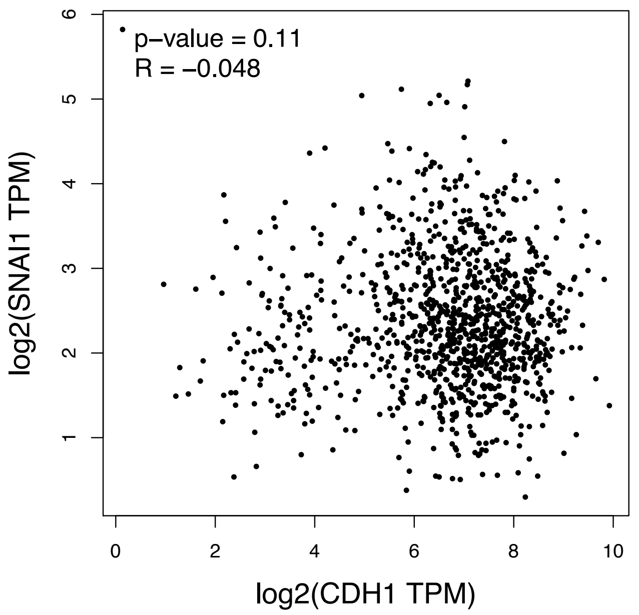


(a)

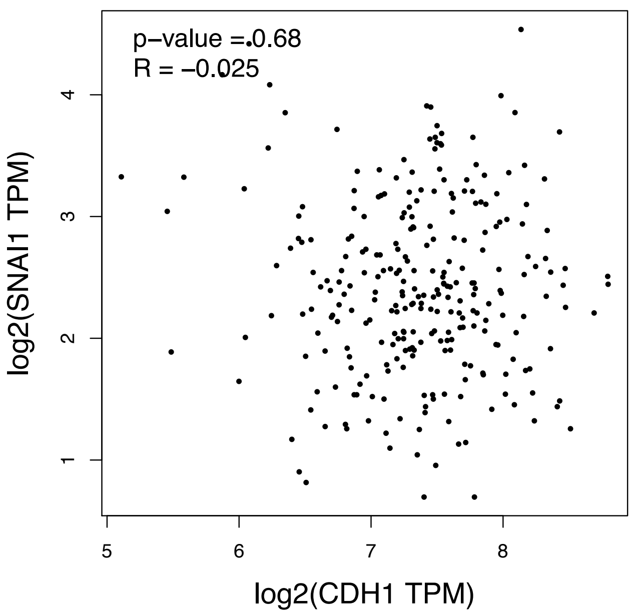


(b)

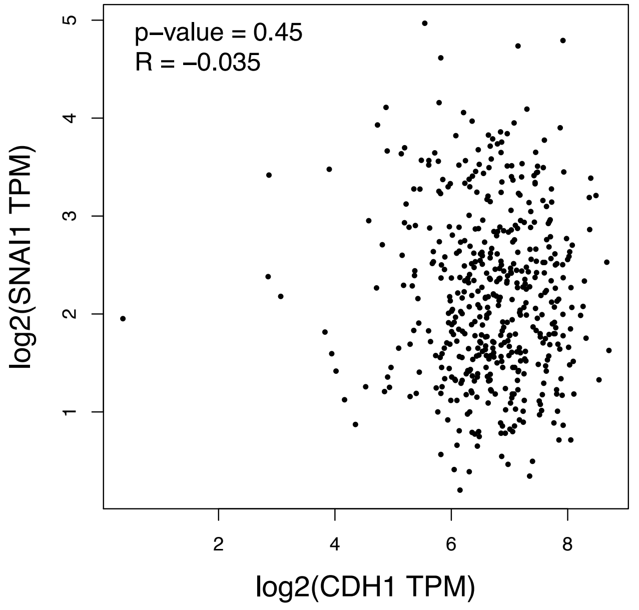


(c)

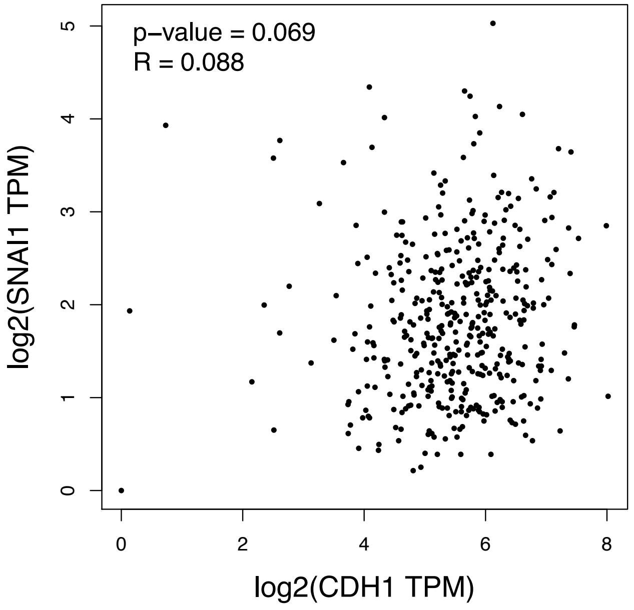


(d)

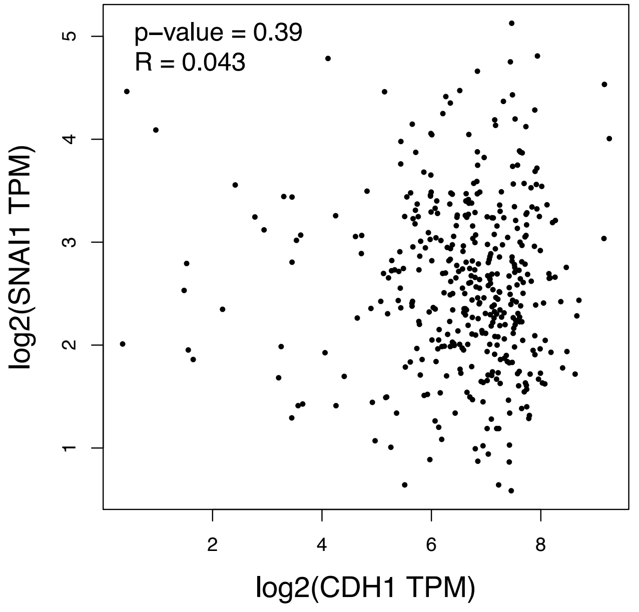


(e)

**Supplementary Figure S3B. Correlation between CDH1 and SNAI1 expression across selected epithelial cancers.**
Pearson correlation analyses between CDH1 and SNAI1 expression were performed using GEPIA2 across **(a)** breast invasive carcinoma (BRCA), **(b)** colon adenocarcinoma (COAD), **(c)** lung adenocarcinoma (LUAD), **(d)** ovarian serous cystadenocarcinoma (OV), and **(e)** stomach adenocarcinoma (STAD). Correlation coefficients and p-values are shown within each plot.

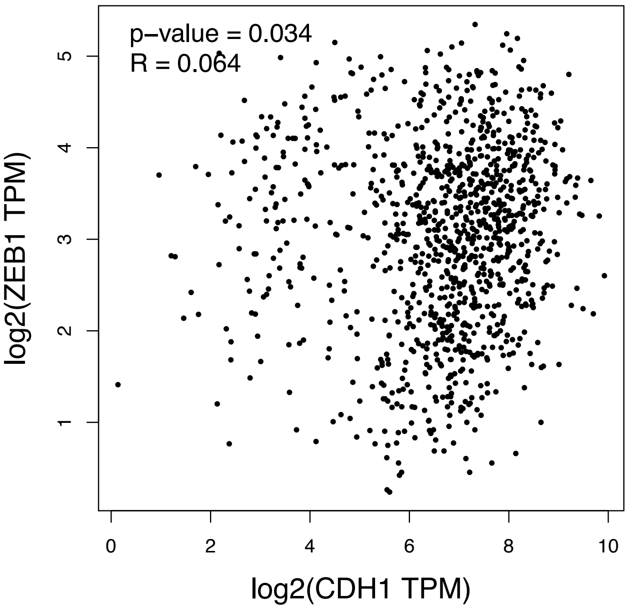


(a)

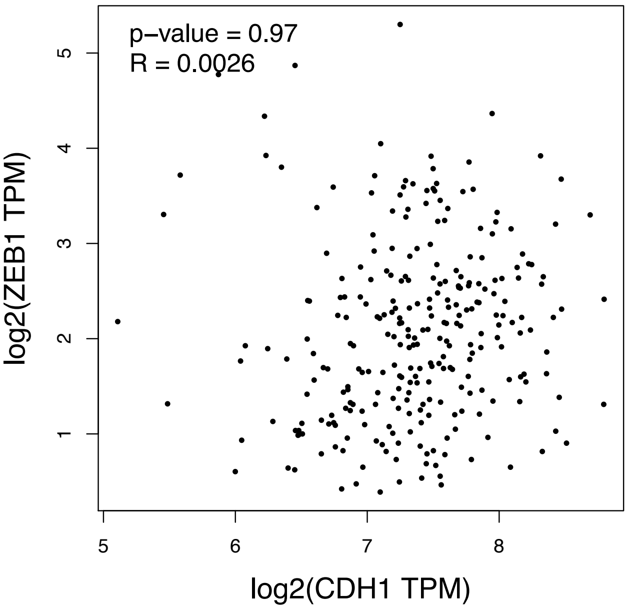


(b)

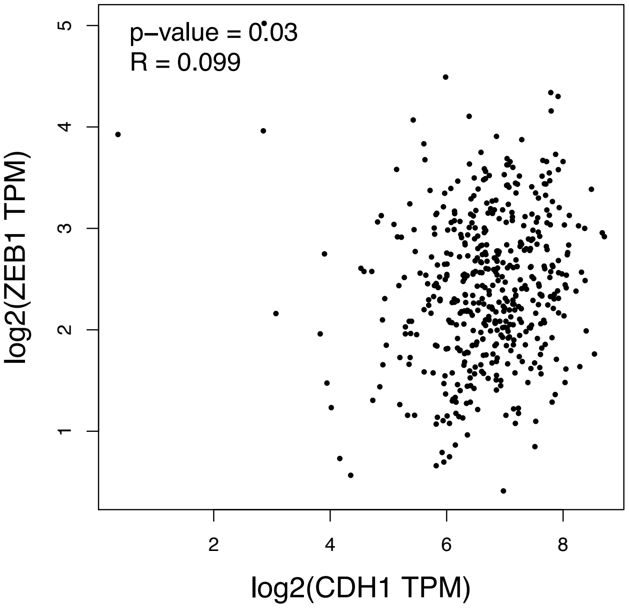


(c)

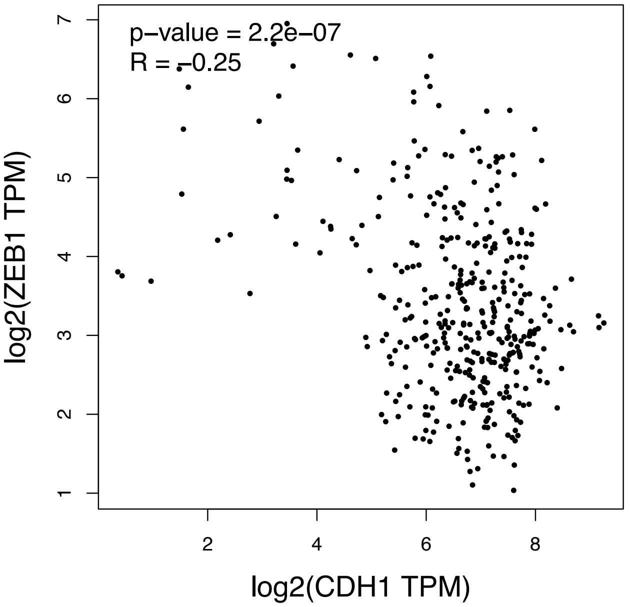


(d)

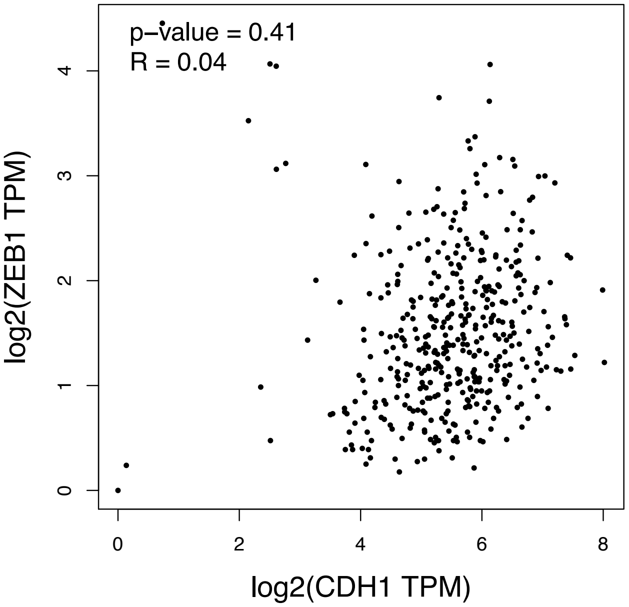


(e)

**Supplementary Figure S3C. Correlation between CDH1 and ZEB1 expression across selected epithelial cancers.**
Pearson correlation analyses between CDH1 and ZEB1 expression were performed using GEPIA2 across (a) breast invasive carcinoma (BRCA), (b) colon adenocarcinoma (COAD), (c) lung adenocarcinoma (LUAD), (d) ovarian serous cystadenocarcinoma (OV), and (e) stomach adenocarcinoma (STAD). Correlation coefficients and p-values are shown within each plot.
