## Supplementary Figure S4 for "Cross-Cancer Profiling of Cadherin-1 Reveals Context-Dependent Epithelial–Mesenchymal Transition Decoupling, Immune Heterogeneity, and Prognostic Variability in Epithelial Cancers"

(a)

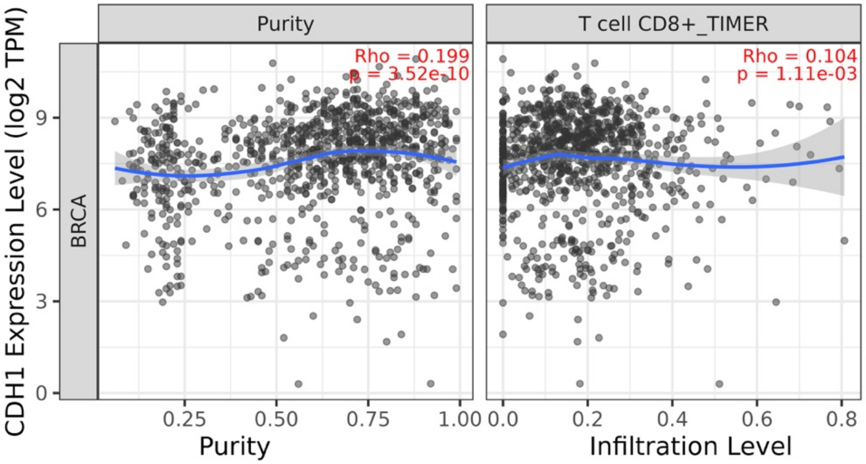


(b)

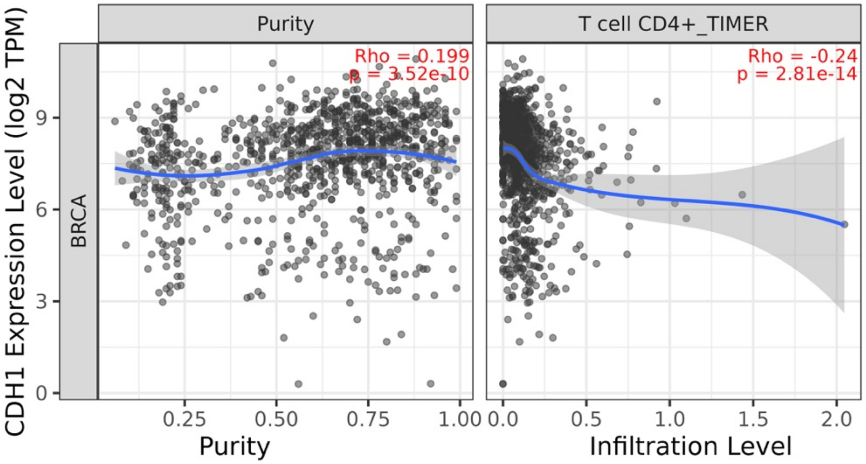


(c)

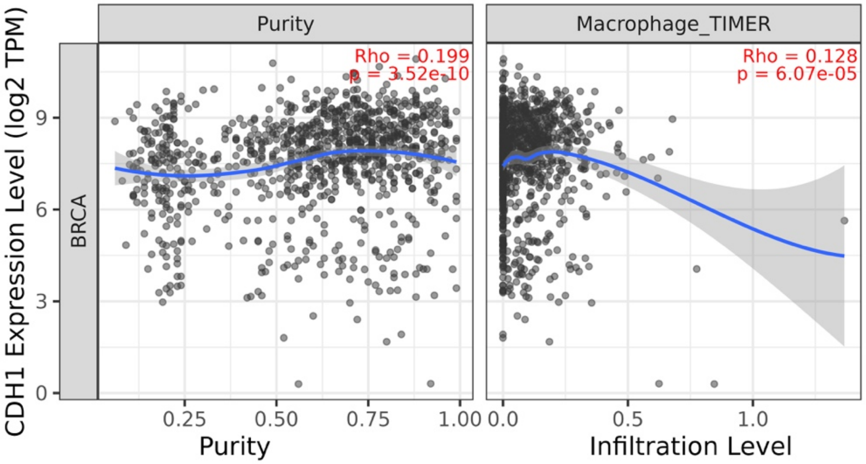


(d)

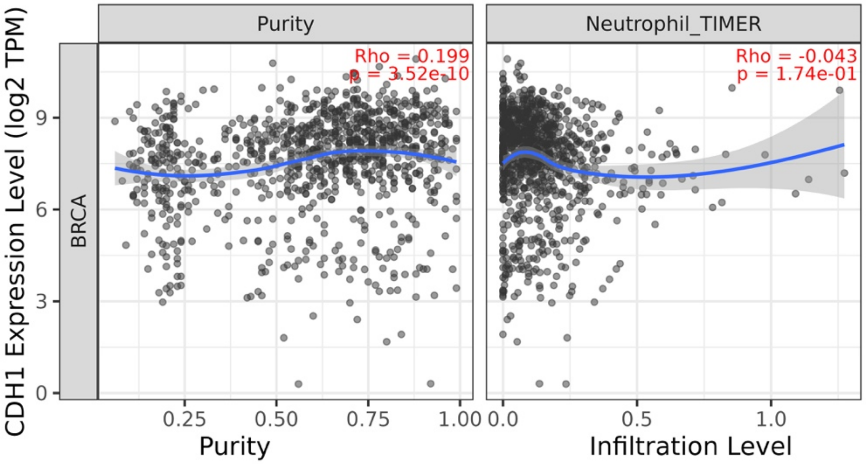


(e)

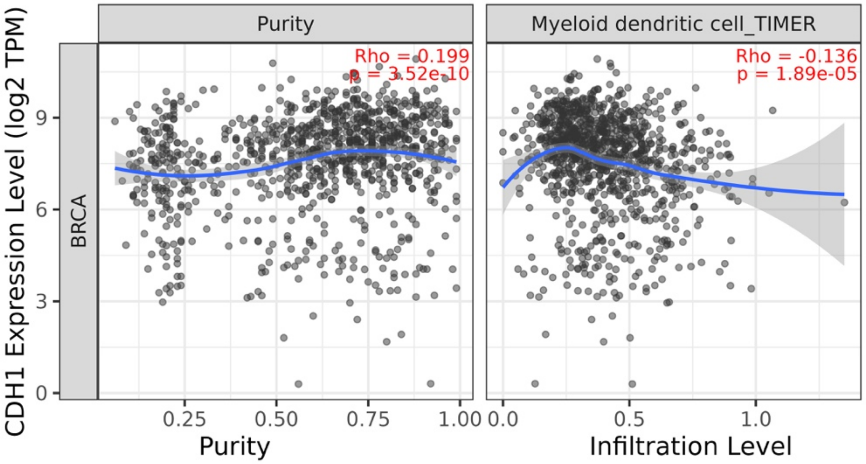


**Supplementary Figure S4A. TIMER3.0 analysis of CDH1-associated immune infiltration in BRCA.**
Purity-adjusted correlations between CDH1 expression and infiltration levels of **(a)** CD8+ T cells, **(b)** CD4+ T cells, **(c)** macrophages, **(d)** neutrophils, and **(e)** dendritic cells in breast invasive carcinoma (BRCA) were analyzed using the TIMER algorithm. For each immune-cell analysis, the left panel shows the relationship between CDH1 expression and tumor purity, while the right panel shows the purity-adjusted association between CDH1 expression and immune-cell infiltration. Correlation coefficients and p-values are shown within each plot.

(a)

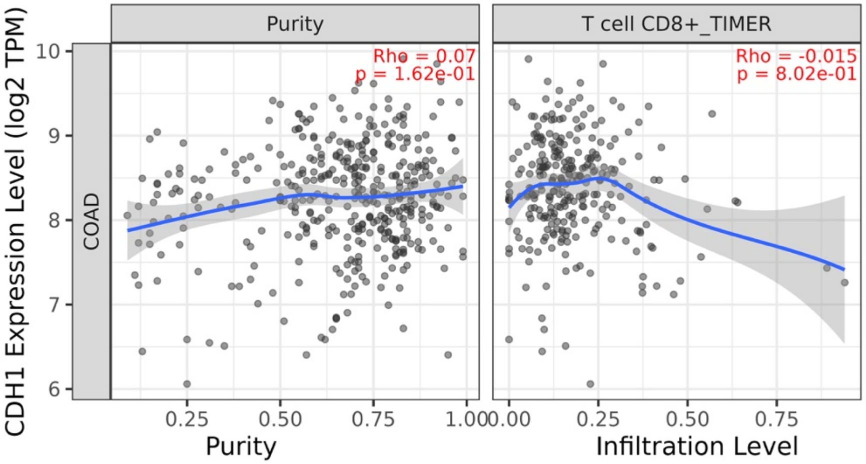


(b)

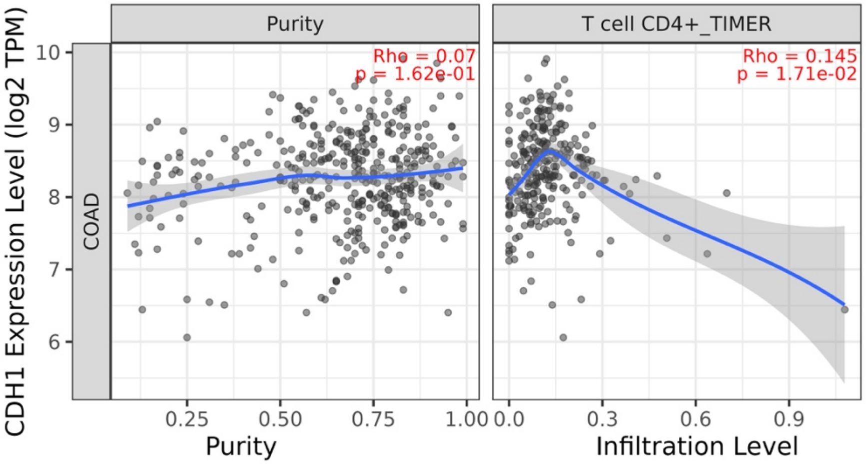


(c)

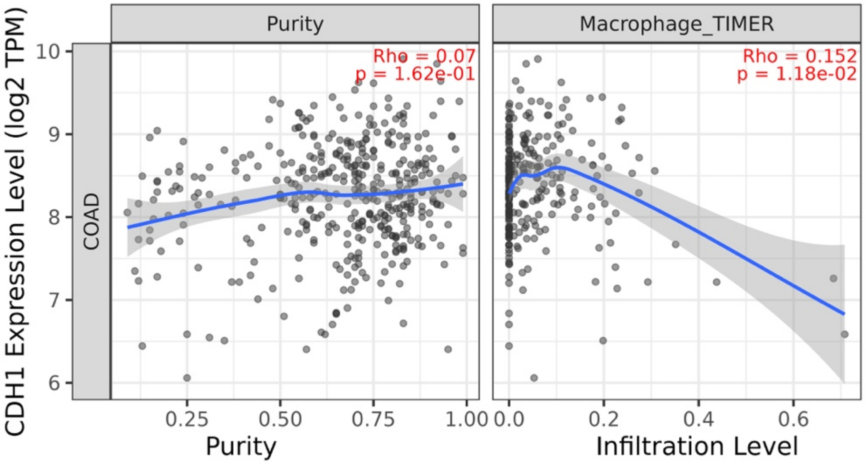


(d)

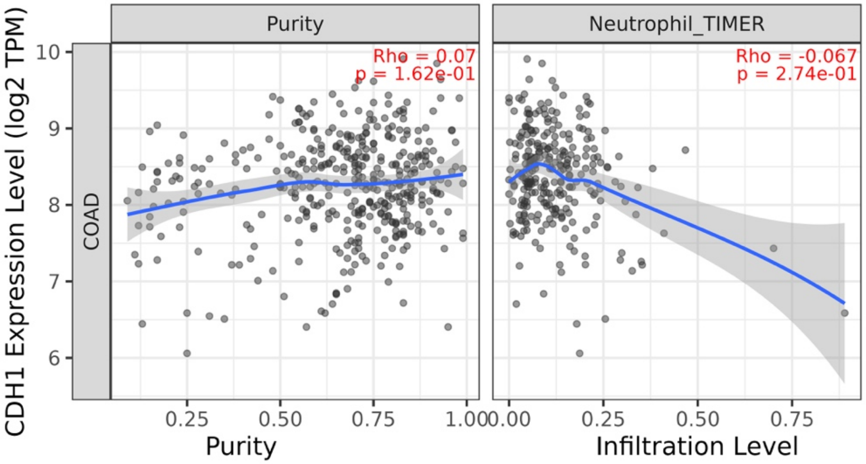


(e)

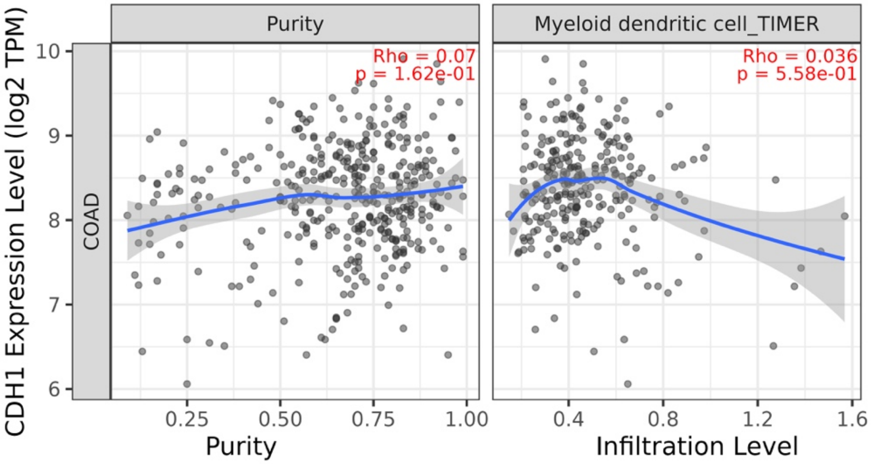


**Supplementary Figure S4B. TIMER3.0 analysis of CDH1-associated immune infiltration in COAD.**
Purity-adjusted correlations between CDH1 expression and infiltration levels of **(a)** CD8+ T cells, **(b)** CD4+ T cells, **(c)** macrophages, **(d)** neutrophils, and **(e)** dendritic cells in colon adenocarcinoma (COAD) were analyzed using the TIMER algorithm. For each immune-cell analysis, the left panel shows the relationship between CDH1 expression and tumor purity, while the right panel shows the purity-adjusted association between CDH1 expression and immune-cell infiltration. Correlation coefficients and p-values are shown within each plot.

(a)

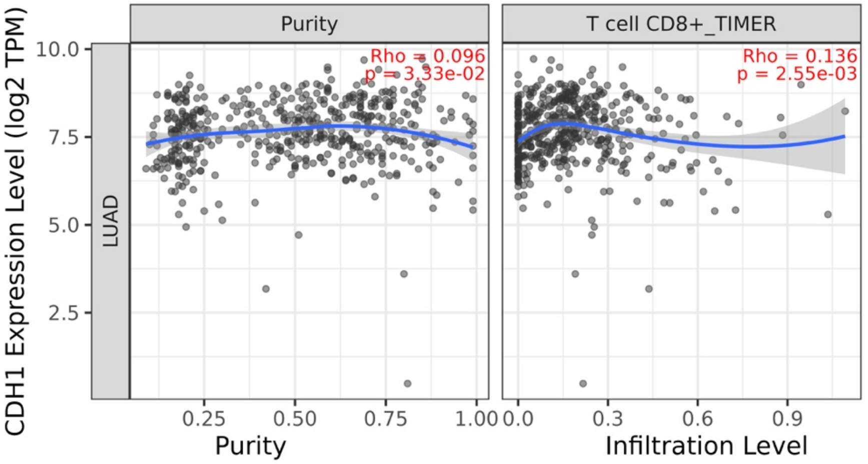


(b)

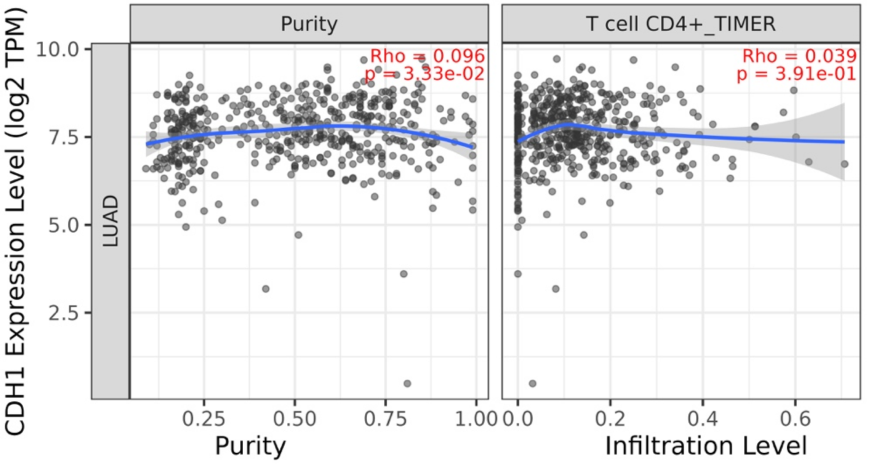


(c)

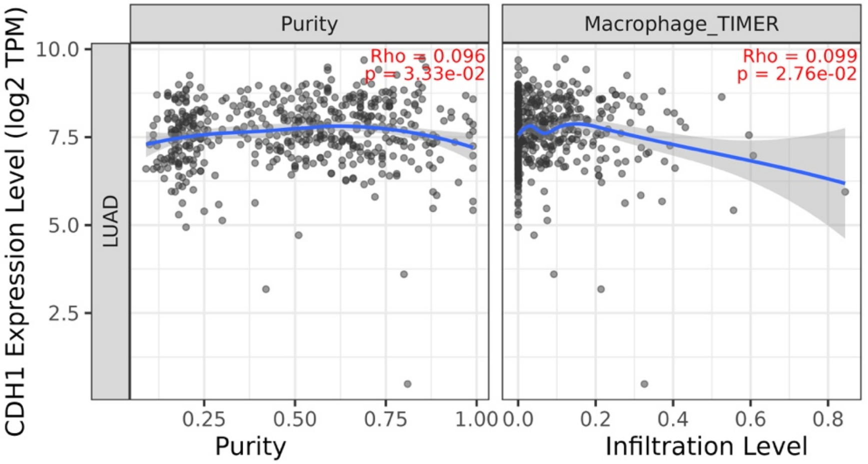


(d)

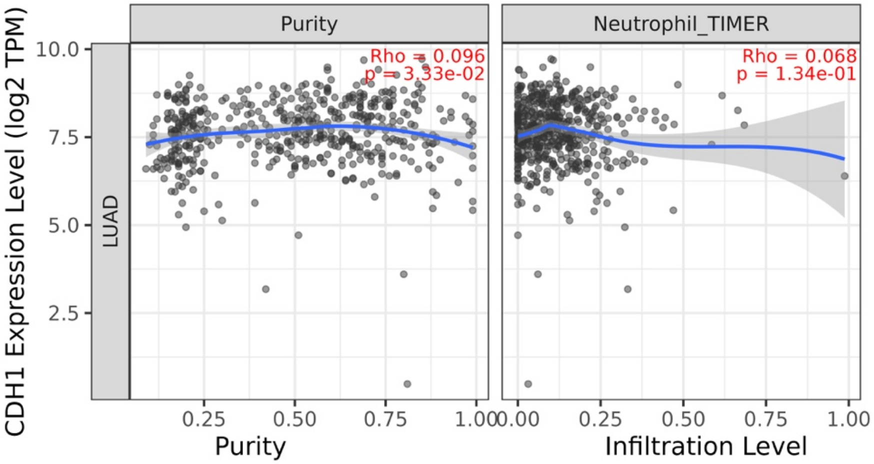


(e)

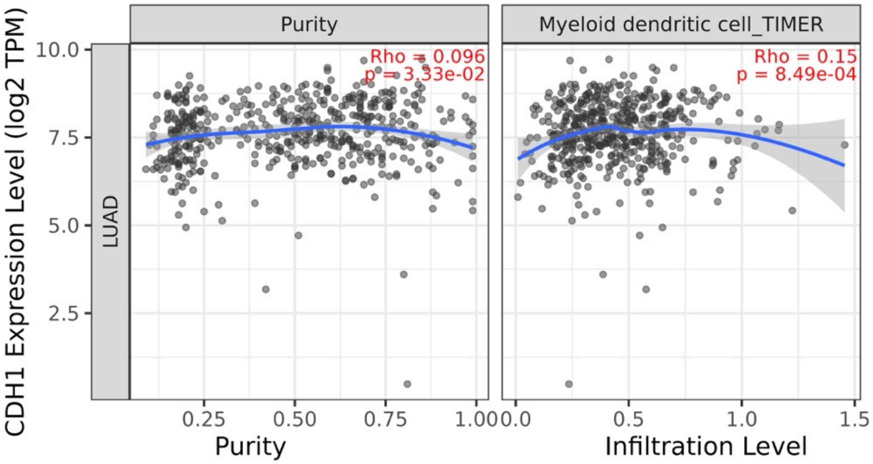


**Supplementary Figure S4C. TIMER3.0 analysis of CDH1-associated immune infiltration in LUAD.**
Purity-adjusted correlations between CDH1 expression and infiltration levels of **(a)** CD8+ T cells, **(b)** CD4+ T cells, **(c)** macrophages, **(d)** neutrophils, and **(e)** dendritic cells in lung adenocarcinoma (LUAD) were analyzed using the TIMER algorithm. For each immune-cell analysis, the left panel shows the relationship between CDH1 expression and tumor purity, while the right panel shows the purity-adjusted association between CDH1 expression and immune-cell infiltration. Correlation coefficients and p-values are shown within each plot.

(a)

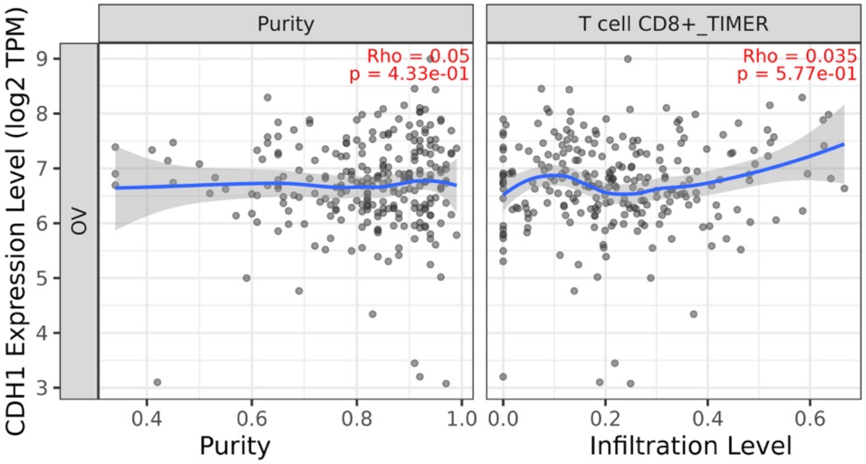


(b)

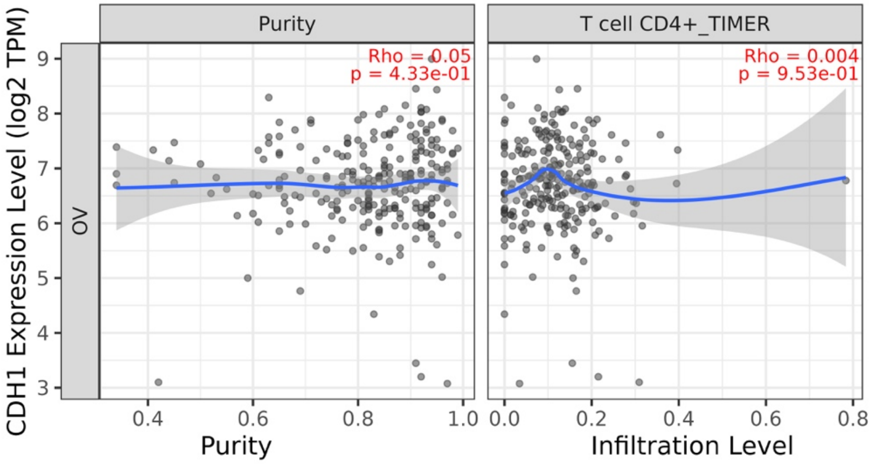


(c)

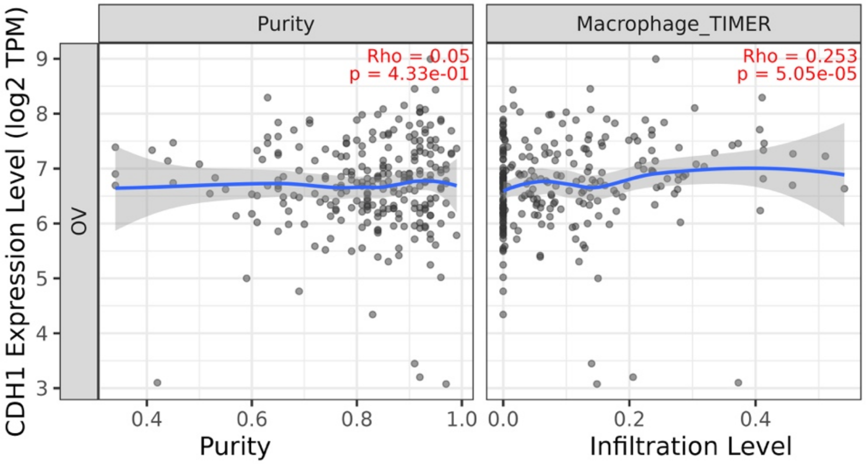


(d)

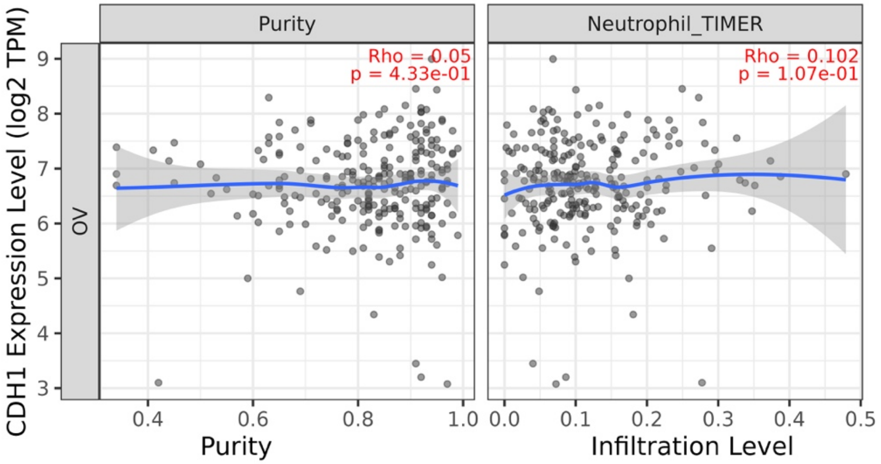


(e)

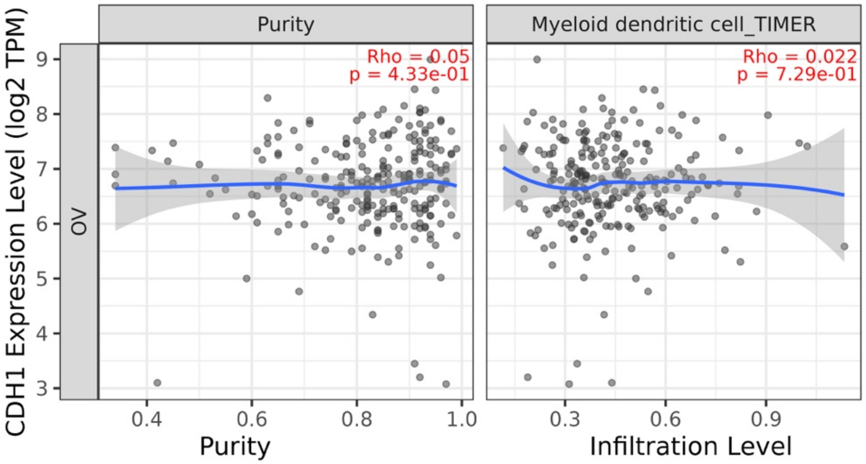


**Supplementary Figure S4D. TIMER3.0 analysis of CDH1-associated immune infiltration in OV.**
Purity-adjusted correlations between CDH1 expression and infiltration levels of **(a)** CD8+ T cells, **(b)** CD4+ T cells, **(c)** macrophages, **(d)** neutrophils, and **(e)** dendritic cells in ovarian serous cystadenocarcinoma (OV) were analyzed using the TIMER algorithm. For each immune-cell analysis, the left panel shows the relationship between CDH1 expression and tumor purity, while the right panel shows the purity-adjusted association between CDH1 expression and immune-cell infiltration. Correlation coefficients and p-values are shown within each plot.

(a)

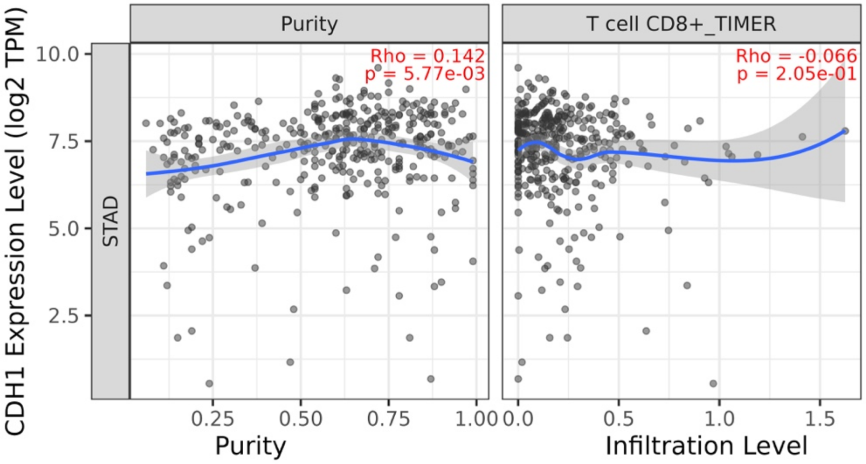


(b)

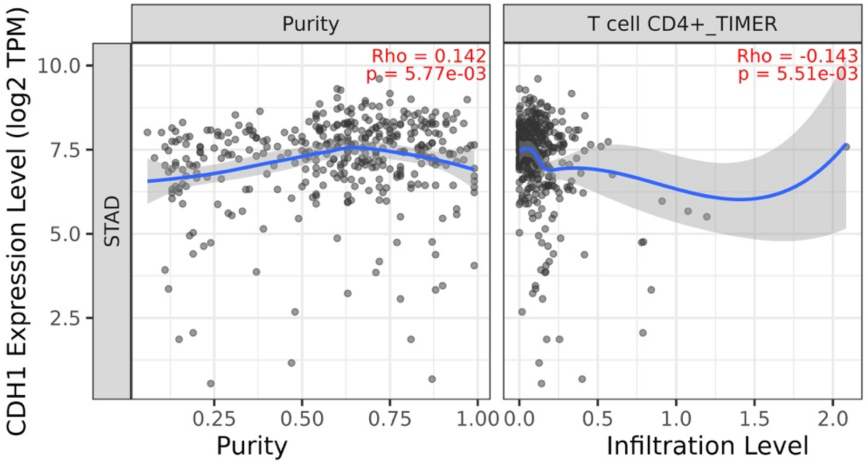


(c)

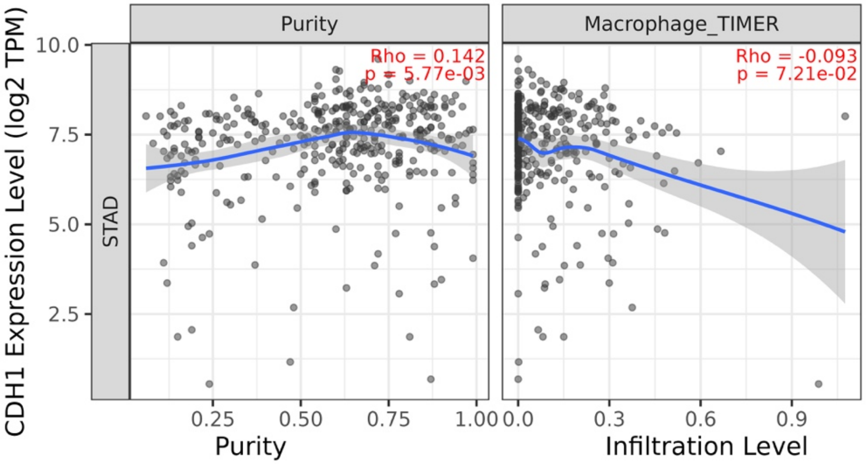


(d)

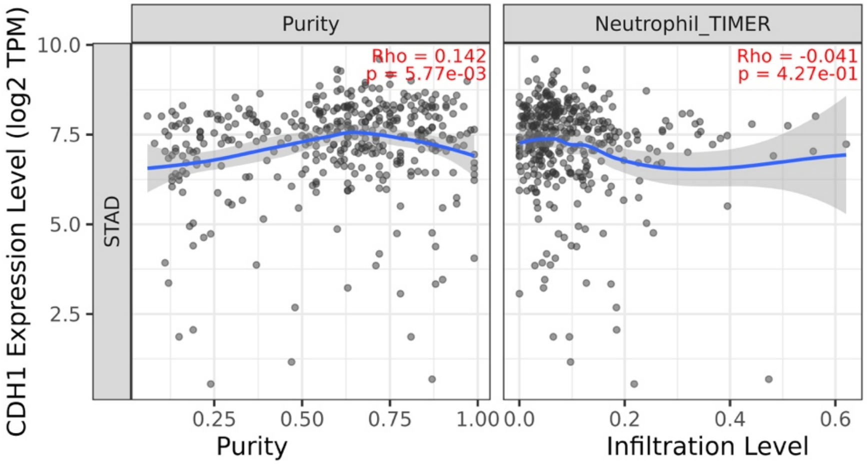


(e)

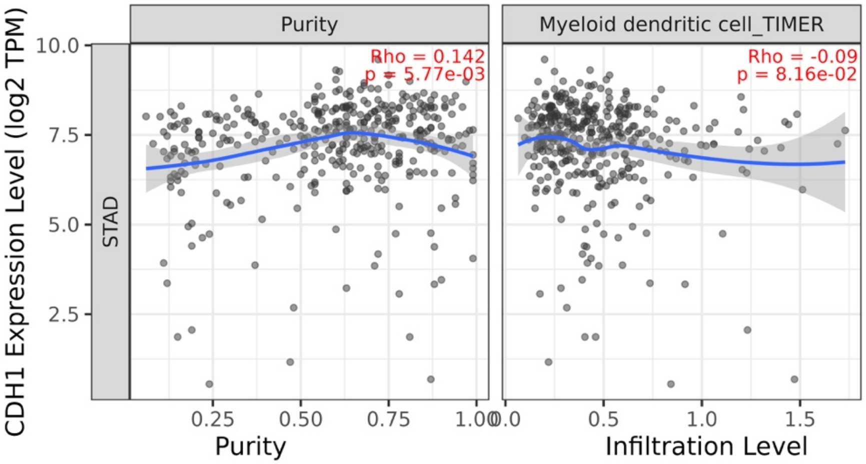


**Supplementary Figure S4E. TIMER3.0 analysis of CDH1-associated immune infiltration in STAD.**
Purity-adjusted correlations between CDH1 expression and infiltration levels of **(a)** CD8+ T cells, **(b)** CD4+ T cells, **(c)** macrophages, **(d)** neutrophils, and **(e)** dendritic cells in stomach adenocarcinoma (STAD) were analyzed using the TIMER algorithm. For each immune-cell analysis, the left panel shows the relationship between CDH1 expression and tumor purity, while the right panel shows the purity-adjusted association between CDH1 expression and immune-cell infiltration. Correlation coefficients and p-values are shown within each plot.
