## Supplementary Figure S5 for "Cross-Cancer Profiling of Cadherin-1 Reveals Context-Dependent Epithelial–Mesenchymal Transition Decoupling, Immune Heterogeneity, and Prognostic Variability in Epithelial Cancers"

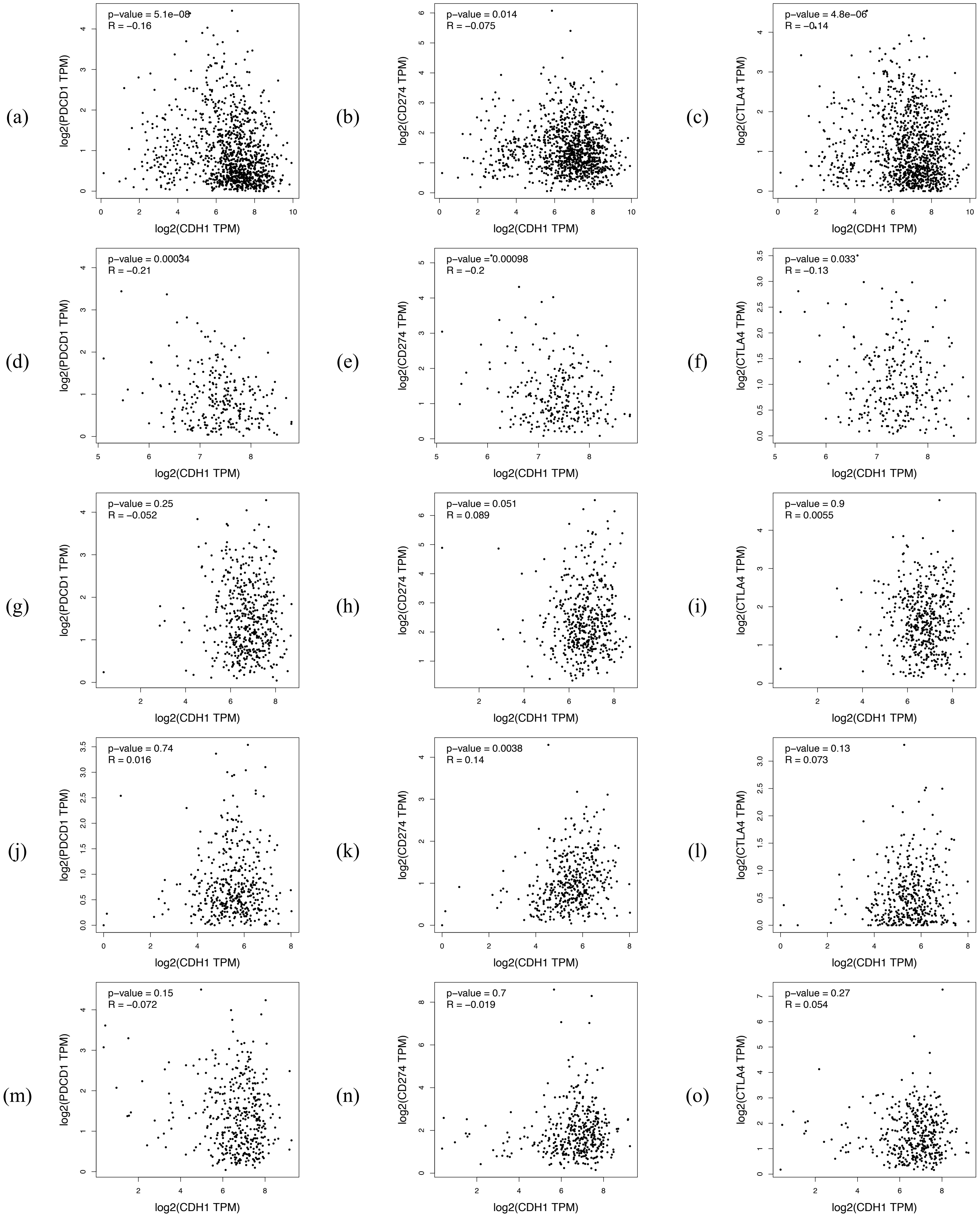

(a)
p−value = 5.1e−08
R = −0.16
log2(PDCD1 TPM)
log2(CDH1 TPM)
(b)
p−value = 0.014
R = −0.075
log2(CD274 TPM)
log2(CDH1 TPM)
(c)
p−value = 4.8e−06
R = −0.14
log2(CTLA4 TPM)
log2(CDH1 TPM)
(d)
p−value = 0.00034
R = −0.21
log2(PDCD1 TPM)
log2(CDH1 TPM)
(e)
p−value = 0.00098
R = −0.2
log2(CD274 TPM)
log2(CDH1 TPM)
(f)
p−value = 0.033
R = −0.13
log2(CTLA4 TPM)
log2(CDH1 TPM)
(g)
p−value = 0.25
R = −0.052
log2(PDCD1 TPM)
log2(CDH1 TPM)
(h)
p−value = 0.051
R = 0.089
log2(CD274 TPM)
log2(CDH1 TPM)
(i)
p−value = 0.9
R = 0.0055
log2(CTLA4 TPM)
log2(CDH1 TPM)
(j)
p−value = 0.74
R = 0.016
log2(PDCD1 TPM)
log2(CDH1 TPM)
(k)
p−value = 0.0038
R = 0.14
log2(CD274 TPM)
log2(CDH1 TPM)
(l)
p−value = 0.13
R = 0.073
log2(CTLA4 TPM)
log2(CDH1 TPM)
(m)
p−value = 0.15
R = −0.072
log2(PDCD1 TPM)
log2(CDH1 TPM)
(n)
p−value = 0.7
R = −0.019
log2(CD274 TPM)
log2(CDH1 TPM)
(o)
p−value = 0.27
R = 0.054
log2(CTLA4 TPM)
log2(CDH1 TPM)

**Supplementary Figure S5. Correlation analysis between CDH1 expression and immune checkpoint genes across selected epithelial cancers.**

Pearson correlation analyses between CDH1 expression and immune checkpoint genes were performed using GEPIA2 across breast invasive carcinoma (BRCA), colon adenocarcinoma (COAD), lung adenocarcinoma (LUAD), ovarian serous cystadenocarcinoma (OV), and stomach adenocarcinoma (STAD). Panels (a–c) show correlations in BRCA, panels (d–f) in COAD, panels (g–i) in LUAD, panels (j–l) in OV, and panels (m–o) in STAD. Within each cancer type, the three panels represent correlations between CDH1 and PDCD1, CD274, and CTLA4, respectively. PDCD1 encodes PD-1, CD274 encodes PD-L1, and CTLA4 encodes CTLA-4. Correlation coefficients and p-values are shown within each plot.
