## Supplementary Figure S6 for "Cross-Cancer Profiling of Cadherin-1 Reveals Context-Dependent Epithelial–Mesenchymal Transition Decoupling, Immune Heterogeneity, and Prognostic Variability in Epithelial Cancers"

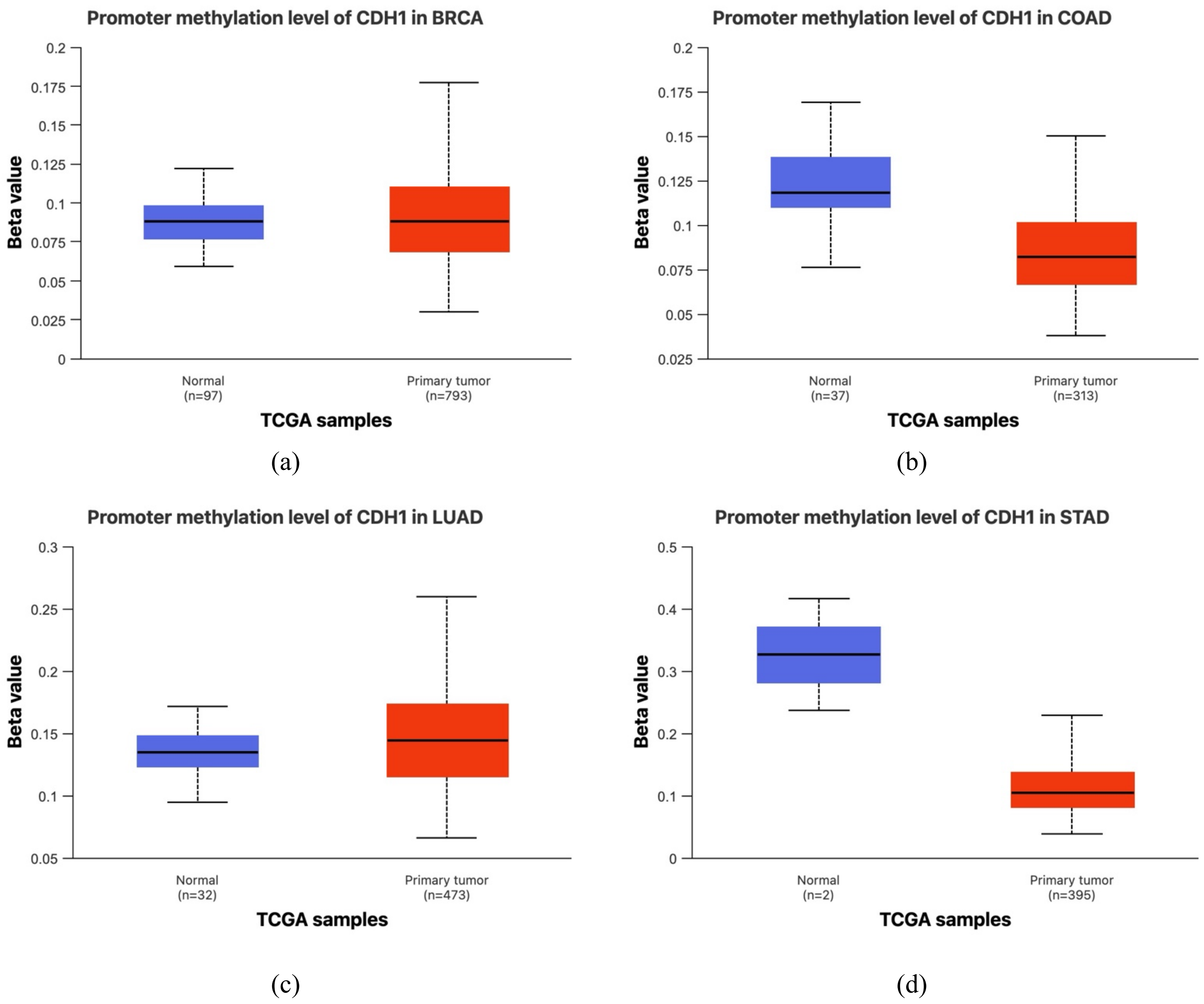


**Supplementary Figure S6. UALCAN analysis of CDH1 promoter methylation across selected epithelial cancers.**

Boxplots showing CDH1 promoter methylation β-values in normal and primary tumor tissues from TCGA datasets for breast invasive carcinoma (BRCA), colon adenocarcinoma (COAD), lung adenocarcinoma (LUAD), and stomach adenocarcinoma (STAD). Ovarian serous cystadenocarcinoma was not included because promoter methylation comparison data were not available in UALCAN. β-values represent DNA methylation levels ranging from 0 to 1. Sample sizes are indicated below each group.
