## Supplementary Figure S7 for "Cross-Cancer Profiling of Cadherin-1 Reveals Context-Dependent Epithelial–Mesenchymal Transition Decoupling, Immune Heterogeneity, and Prognostic Variability in Epithelial Cancers"

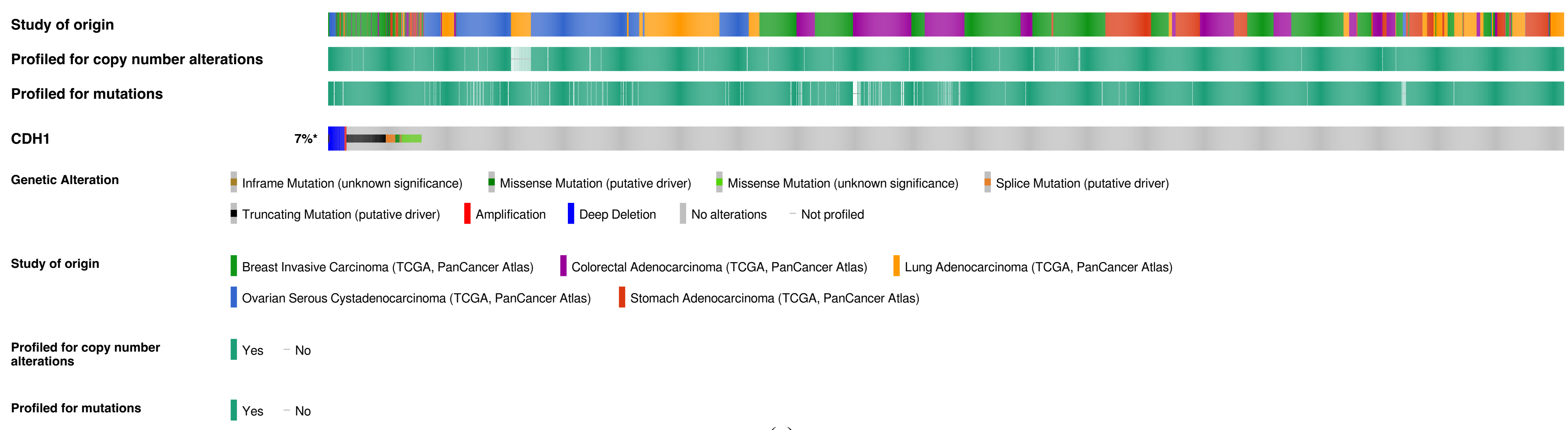
Study of origin
Profiled for copy number alterations
Profiled for mutations
CDH1
7%*
Genetic Alteration
Inframe Mutation (unknown significance)
Missense Mutation (putative driver)
Missense Mutation (unknown significance)
Splice Mutation (putative driver)
Truncating Mutation (putative driver)
Amplification
Deep Deletion
No alterations
Not profiled
Study of origin
Breast Invasive Carcinoma (TCGA, PanCancer Atlas)
Colorectal Adenocarcinoma (TCGA, PanCancer Atlas)
Lung Adenocarcinoma (TCGA, PanCancer Atlas)
Ovarian Serous Cystadenocarcinoma (TCGA, PanCancer Atlas)
Stomach Adenocarcinoma (TCGA, PanCancer Atlas)
Profiled for copy number alterations
Yes
No
Profiled for mutations
Yes
No

(a)

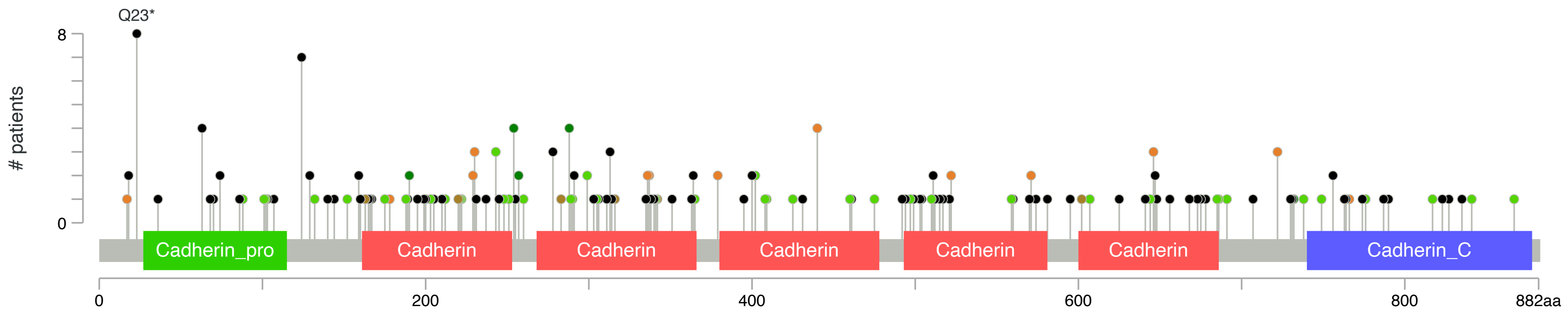
Q23*
8
0
### patients
Cadherin_pro
Cadherin
Cadherin
Cadherin
Cadherin
Cadherin
Cadherin_C
0
200
400
600
800
882aa

(b)

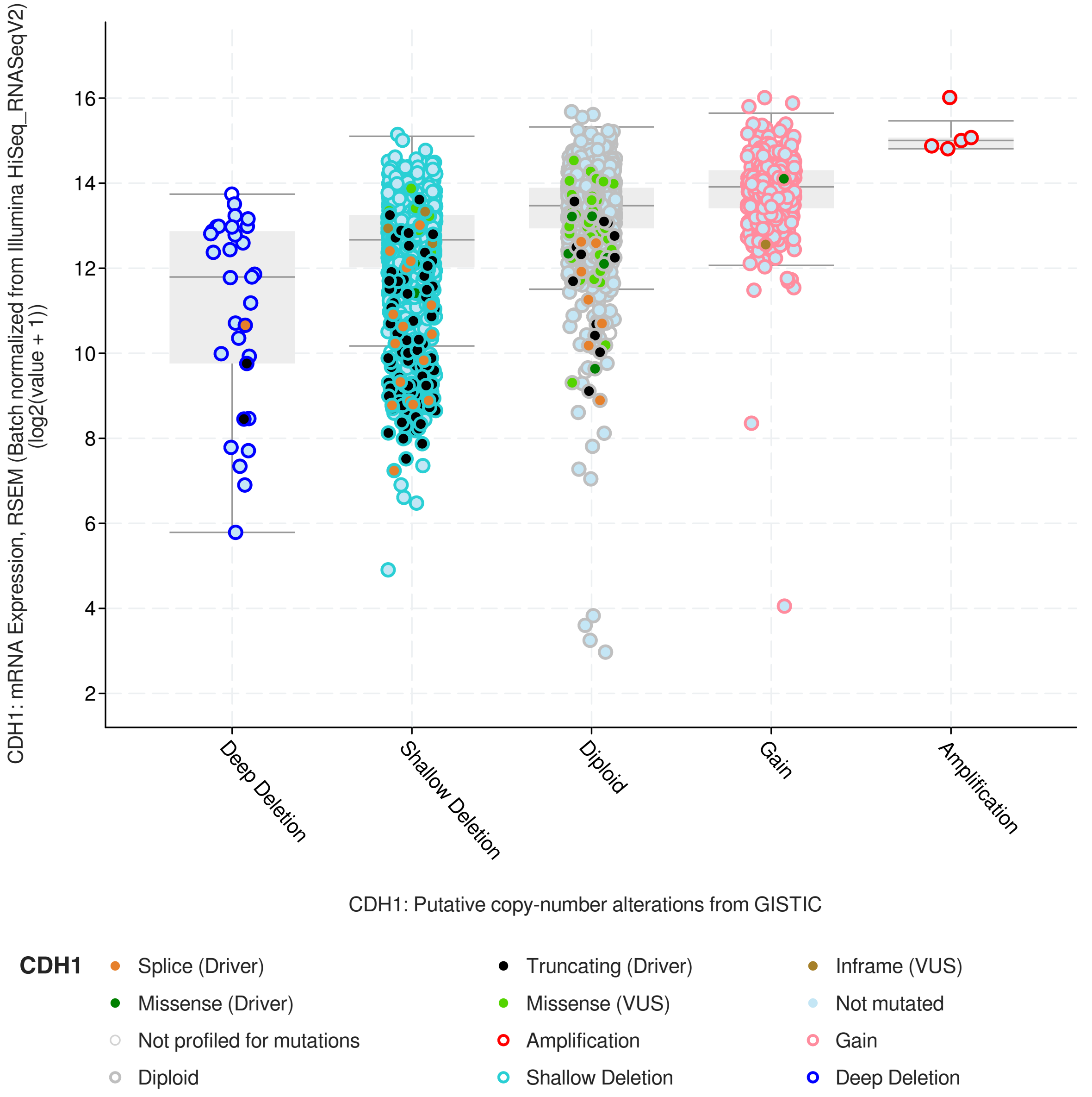


(c)

**Supplementary Figure S7. cBioPortal analysis of CDH1 genomic alterations across selected epithelial cancers.**
(a) OncoPrint showing sample-level CDH1 genomic alterations across breast invasive carcinoma (BRCA), colon adenocarcinoma (COAD), lung adenocarcinoma (LUAD), ovarian serous cystadenocarcinoma (OV), and stomach adenocarcinoma (STAD) from TCGA PanCancer Atlas cohorts. Alteration types include somatic mutations and copy-number alterations as annotated by cBioPortal. (b) Lollipop plot showing the distribution of CDH1 mutations across the encoded E-cadherin protein sequence. Mutation positions are mapped along annotated protein domains where available. (c) Association between CDH1 copy-number alteration status and mRNA expression. Copy-number categories were derived from GISTIC calls and include deep deletion, shallow deletion, diploid, gain, and amplification where available. CNA, copy-number alteration; TCGA, The Cancer Genome Atlas; GISTIC, Genomic Identification of Significant Targets in Cancer.
