## Supplementary Figure S8 for "Cross-Cancer Profiling of Cadherin-1 Reveals Context-Dependent Epithelial–Mesenchymal Transition Decoupling, Immune Heterogeneity, and Prognostic Variability in Epithelial Cancers"

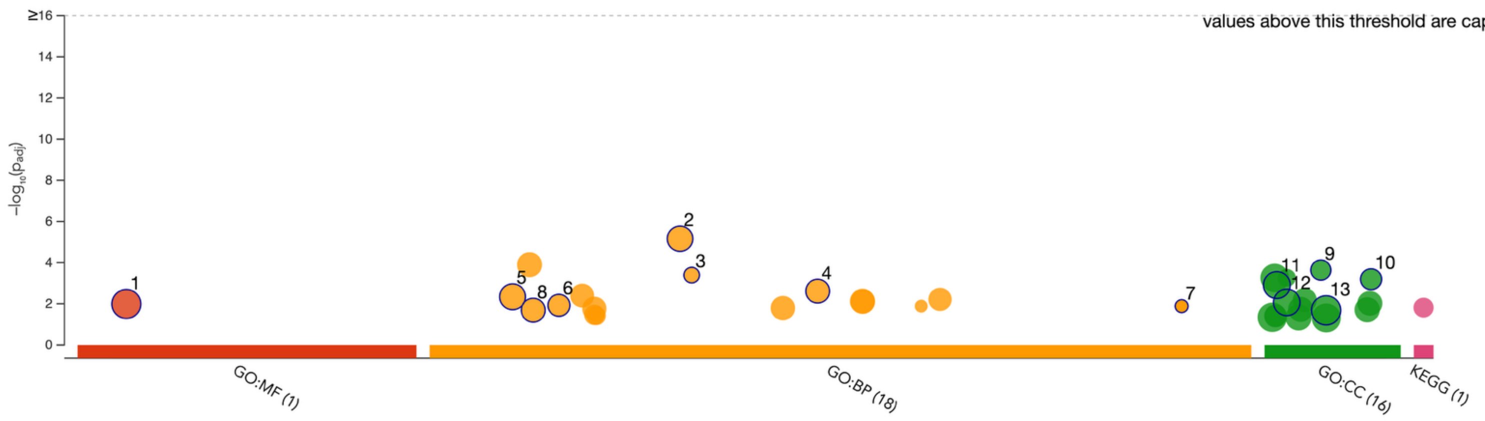


(a)

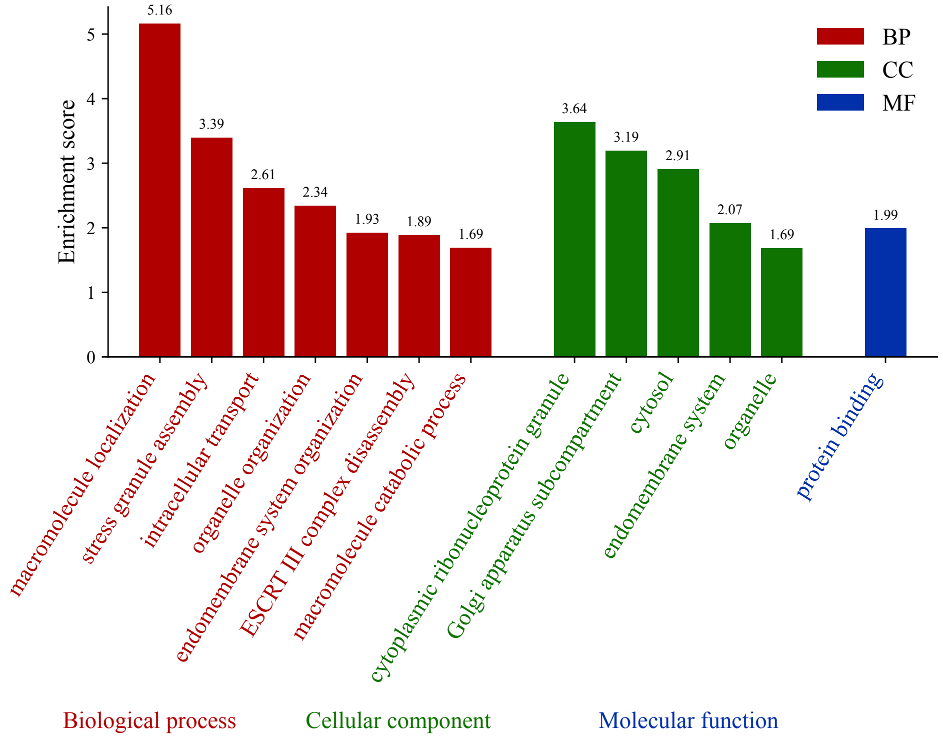


(b)

**Supplementary Figure S8. Functional enrichment analysis of CDH1-correlated genes.**
(a) g:Profiler enrichment overview showing enriched Gene Ontology and KEGG terms among the top 100 genes positively correlated with CDH1 identified using GEPIA2. (b) Bar plot of selected enriched Gene Ontology terms grouped by biological process, cellular component, and molecular function. Enrichment score represents −log10 adjusted p-value, with higher values indicating stronger enrichment. BP, biological process; CC, cellular component; MF, molecular function; GO, Gene Ontology; KEGG, Kyoto Encyclopedia of Genes and Genomes.
