## Supplementary Figure S9 for "Cross-Cancer Profiling of Cadherin-1 Reveals Context-Dependent Epithelial–Mesenchymal Transition Decoupling, Immune Heterogeneity, and Prognostic Variability in Epithelial Cancers"

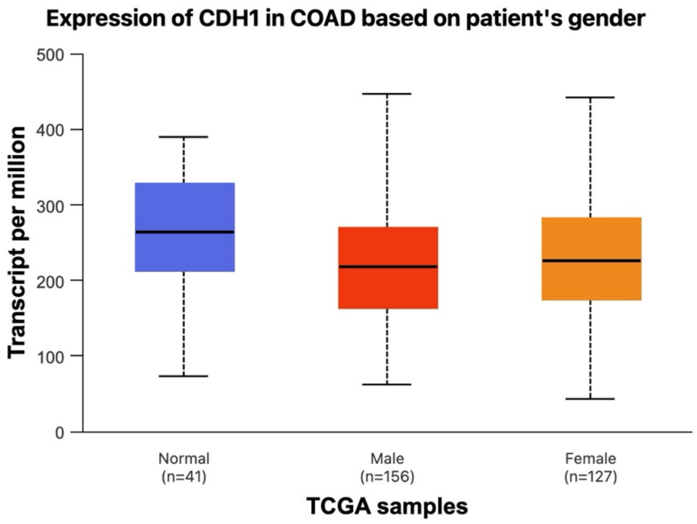


(a)

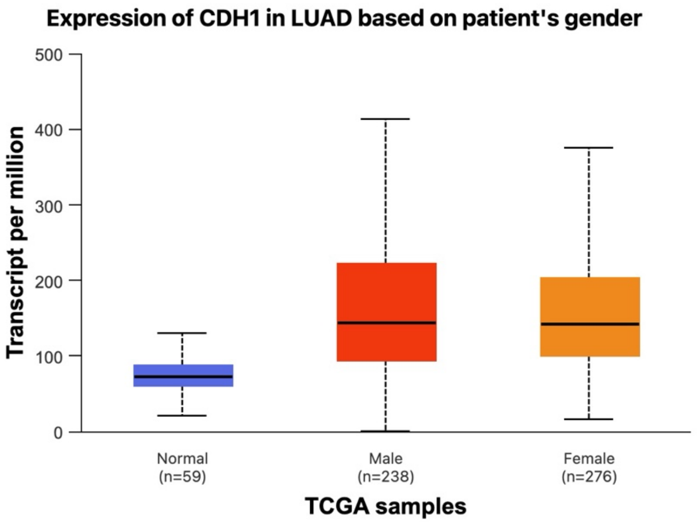


(b)

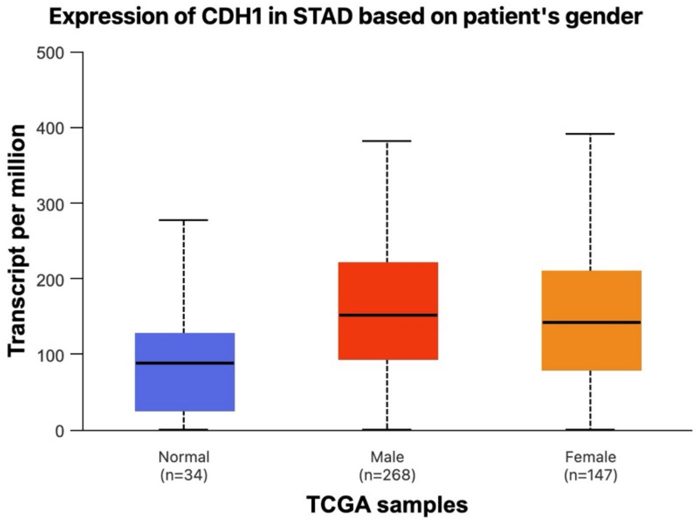


(c)

**Supplementary Figure S9. Sex-stratified UALCAN analysis of CDH1 expression in selected epithelial cancers.**
Boxplots showing CDH1 transcript expression according to patient sex in (a) colon adenocarcinoma (COAD), (b) lung adenocarcinoma (LUAD), and (c) stomach adenocarcinoma (STAD) using UALCAN TCGA datasets. Normal tissue groups are shown for comparison where available. Tumor samples were stratified by male and female patient groups. Sample sizes are indicated below each group.
