## Supplementary Table S1 for "Cross-Cancer Profiling of Cadherin-1 Reveals Context-Dependent Epithelial–Mesenchymal Transition Decoupling, Immune Heterogeneity, and Prognostic Variability in Epithelial Cancers"

**Supplementary Table S1. TCGA-based overall survival analysis according to CDH1 expression**

| **Cancer** | **Analyzed cases (n)** | **Cutoff ^a^** | **HR** | **95% CI** | **p-value** | **Median OS, low CDH1 (days)** | **Median OS, high CDH1 (days)** |
| --- | --- | --- | --- | --- | --- | --- | --- |
| BRCA | 1203 | 7.87 | 1.00 | 0.75–1.32 | 0.980 | 3959 | 3461 |
| LUAD | 589 | 13.96 | 0.98 | 0.75–1.29 | 0.910 | 1379 | 1528 |
| COAD | 488 | 8.29 | 0.80 | 0.55–1.17 | 0.240 | 2821 | 2532 |
| OV | 428 | 6.58 | 1.09 | 0.86–1.39 | 0.480 | 1366 | 1354 |
| STAD | 420 | 7.37 | 1.00 | 0.74–1.35 | 0.990 | 1095 | 874 |

^a^ Cutoff values represent median CDH1 expression thresholds used to stratify patients into low- and high-expression groups.
BRCA, breast invasive carcinoma; COAD, colon adenocarcinoma; LUAD, lung adenocarcinoma; OV, ovarian serous cystadenocarcinoma; STAD, stomach adenocarcinoma; TCGA, The Cancer Genome Atlas; OS, overall survival; HR, hazard ratio; CI, confidence interval.
