## Supplementary Table S2 for "Cross-Cancer Profiling of Cadherin-1 Reveals Context-Dependent Epithelial–Mesenchymal Transition Decoupling, Immune Heterogeneity, and Prognostic Variability in Epithelial Cancers"

**Supplementary Table S2. Pearson correlations between CDH1 and EMT-associated genes across selected epithelial cancers**

| **Cancer** | **VIM**  **(R, p-value)** | **SNAI1**  **(R, p-value)** | **ZEB1**  **(R, p-value)** | **Interpretation** |
| --- | --- | --- | --- | --- |
| BRCA | -0.20, <0.001 | -0.048, 0.110 | 0.064, 0.034 | Non-canonical EMT |
| COAD | -0.12, 0.045 | -0.025, 0.680 | 0.0026, 0.970 | Weak EMT association |
| LUAD | 0.0017, 0.970 | -0.035, 0.450 | 0.099, 0.030 | No classical EMT |
| OV | 0.08, 0.100 | 0.088, 0.069 | 0.04, 0.410 | No significant EMT |
| STAD | -0.27, <0.001 | 0.043, 0.390 | -0.25, <0.001 | Classical EMT pattern |

BRCA, breast invasive carcinoma; COAD, colon adenocarcinoma; LUAD, lung adenocarcinoma; OV, ovarian serous cystadenocarcinoma; STAD, stomach adenocarcinoma; EMT, epithelial–mesenchymal transition; R, Pearson correlation coefficient.
