## Supplementary Table S3 for "Cross-Cancer Profiling of Cadherin-1 Reveals Context-Dependent Epithelial–Mesenchymal Transition Decoupling, Immune Heterogeneity, and Prognostic Variability in Epithelial Cancers"

**Supplementary Table S3. TIMER-derived correlations between CDH1 expression and immune cell infiltration**

| **Cancer** | **CD8+ T cells (ρ, p)** | **CD4+ T cells (ρ, p)** | **Macrophages (ρ, p)** | **Neutrophils  (ρ, p)** | **Dendritic cells (ρ, p)** | **Interpretation** |
| --- | --- | --- | --- | --- | --- | --- |
| BRCA | 0.104, 0.001 | -0.240, < 0.001 | 0.128, < 0.001 | -0.043, 0.174 | -0.136, < 0.001 | Mixed immune modulation |
| COAD | -0.015, 0.802 | 0.145, 0.017 | 0.152, 0.012 | -0.067, 0.274 | 0.036, 0.558 | Helper/macrophage-skewed |
| LUAD | 0.136, 0.003 | 0.039, 0.391 | 0.099, 0.028 | 0.068, 0.134 | 0.150, < 0.001 | Cytotoxic + APC involvement |
| OV | 0.035, 0.577 | 0.004, 0.953 | 0.253, < 0.001 | 0.102, 0.107 | 0.022, 0.729 | Macrophage-dominant |
| STAD | -0.066, 0.205 | -0.143, 0.006 | -0.093, 0.072 | -0.041, 0.427 | -0.090, 0.082 | Limited immune association |

APC, antigen-presenting cell; BRCA, breast invasive carcinoma; COAD, colon adenocarcinoma; LUAD, lung adenocarcinoma; OV, ovarian serous cystadenocarcinoma; STAD, stomach adenocarcinoma; TIMER, Tumor Immune Estimation Resource; ρ, correlation coefficient.
