## Supplementary Table S4 for "Cross-Cancer Profiling of Cadherin-1 Reveals Context-Dependent Epithelial–Mesenchymal Transition Decoupling, Immune Heterogeneity, and Prognostic Variability in Epithelial Cancers"

**Supplementary Table S4. Correlations between CDH1 expression and immune checkpoint genes**

| **Cancer** | **PDCD1 (R, p)** | **CD274 (R, p)** | **CTLA4 (R, p)** | **Interpretation** |
| --- | --- | --- | --- | --- |
| BRCA | -0.160, < 0.001 | -0.075, 0.014 | -0.140, < 0.001 | Weak inverse association |
| COAD | -0.210, < 0.001 | -0.200, < 0.001 | -0.130, 0.033 | Consistent inverse association |
| LUAD | -0.052, 0.250 | 0.089, 0.051 | 0.006, 0.900 | No association |
| OV | 0.016, 0.740 | 0.140, 0.004 | 0.073, 0.130 | CD274-specific positive association |
| STAD | -0.072, 0.150 | -0.019, 0.700 | 0.054, 0.270 | No association |

BRCA, breast invasive carcinoma; COAD, colon adenocarcinoma; LUAD, lung adenocarcinoma; OV, ovarian serous cystadenocarcinoma; R, Pearson correlation coefficient; STAD, stomach adenocarcinoma.
