## Supplementary Table S5 for "Cross-Cancer Profiling of Cadherin-1 Reveals Context-Dependent Epithelial–Mesenchymal Transition Decoupling, Immune Heterogeneity, and Prognostic Variability in Epithelial Cancers"

**Supplementary Table S5. cBioPortal-derived CDH1 genomic alteration frequencies across selected epithelial cancers**

| **Cancer** | **Alteration frequency (%)** | **Mutation (%)** | **Amplification (%)** | **Deep deletion (%)** | **Multiple Alterations (%)** | **Dominant alteration** | **Interpretation** |
| --- | --- | --- | --- | --- | --- | --- | --- |
| BRCA | 13.65 (148/1084) | 11.72 | 0.28 | 1.38 | 0.28 | Mutation | Moderate alteration; mutation-driven subset present |
| COAD | 3.37 (20/594) | 3.20 | - | 0.17 | - | Mutation | Low-frequency mutation; minimal CNA |
| LUAD | 1.41 (8/566) | 1.06 | 0.18 | 0.18 | - | Mutation (very low) | Minimal genomic involvement |
| OV | 3.77 (22/584) | 0.68 | - | 3.08 | - | Deep deletion | Low mutation; deletion-driven pattern |
| STAD | 10.68 (47/440) | 9.32 | 0.23 | 1.14 | - | Mutation | Mutation contributes in subset; not dominant overall |

Alteration frequencies were obtained from cBioPortal using TCGA PanCancer Atlas cohorts. Values in parentheses indicate altered cases/total profiled cases.
BRCA, breast invasive carcinoma; CNA, copy-number alteration; COAD, colon adenocarcinoma; LUAD, lung adenocarcinoma; OV, ovarian serous cystadenocarcinoma; STAD, stomach adenocarcinoma; TCGA, The Cancer Genome Atlas.
- indicates that the alteration type was not detected or not reported for that cohort.
