## Supplementary Table S6 for "Cross-Cancer Profiling of Cadherin-1 Reveals Context-Dependent Epithelial–Mesenchymal Transition Decoupling, Immune Heterogeneity, and Prognostic Variability in Epithelial Cancers"

**Supplementary Table S6. Sex-stratified UALCAN analysis of CDH1 expression across epithelial cancers**

| **Cancer** | **Normal vs Male  (p-value)** | **Normal vs Female  (p-value)** | **Male vs Female  (p-value)** | **Overall interpretation** |
| --- | --- | --- | --- | --- |
| COAD | 0.105 | 0.122 | 0.794 | No significant sex-dependent differences |
| LUAD | < 0.001 | < 0.001 | 0.777 | Tumor-associated upregulation independent of sex |
| STAD | < 0.001 | < 0.001 | 0.122 | Tumor-associated upregulation independent of sex |

P-values were obtained from UALCAN sex-stratified expression analysis comparing normal tissues, male tumor samples, and female tumor samples.
COAD, colon adenocarcinoma; LUAD, lung adenocarcinoma; STAD, stomach adenocarcinoma
